# Quantitative proteomics of the 2016 WHO *Neisseria gonorrhoeae* reference strains surveys vaccine candidates and antimicrobial resistance determinants

**DOI:** 10.1101/434753

**Authors:** Fadi E. El-Rami, Ryszard A. Zielke, Teodora Wi, Aleksandra E. Sikora, Magnus Unemo

## Abstract

The sexually transmitted disease gonorrhea (causative agent: *Neisseria gonorrhoeae*) remains an urgent public health threat globally due to the repercussions on reproductive health, high incidence, widespread antimicrobial resistance (AMR), and absence of a vaccine. To mine gonorrhea antigens and enhance our understanding of gonococcal AMR at the proteome level, we performed the first large-scale proteomic profiling of a diverse panel (*n*=15) of gonococcal strains, including the 2016 World Health Organization (WHO) reference strains. These strains show all existing AMR profiles, previously described in regard to phenotypic and reference genome characteristics, and are intended for quality assurance in laboratory investigations. Herein, these isolates were subjected to subcellular fractionation and labeling with tandem mass tags coupled to mass spectrometry and multi-combinatorial bioinformatics. Our analyses detected 901 and 723 common proteins in cell envelope and cytoplasmic subproteomes, respectively. We identified nine novel gonorrhea vaccine candidates. Expression and conservation of new and previously selected antigens were investigated. In addition, established gonococcal AMR determinants were evaluated for the first time using quantitative proteomics. Six new proteins, WHO_F_00238, WHO_F_00635, WHO_F_00745, WHO_F_01139, WHO_F_01144, and WHO_F_01226, were differentially expressed in all strains, suggesting that they represent global proteomic AMR markers, indicate a predisposition toward developing or compensating gonococcal AMR, and/or act as new antimicrobial targets. Finally, phenotypic clustering based on the isolates’ defined antibiograms and common differentially expressed proteins yielded seven matching clusters between established and proteome-derived AMR signatures. Together, our investigations provide a reference proteomics databank for gonococcal vaccine and AMR research endeavors, which enables microbiological, clinical, or epidemiological projects and enhances the utility of the WHO reference strains.

## INTRODUCTION

*Neisseria gonorrhoeae* is an obligate human pathogen and the causative agent of the sexually transmitted disease gonorrhea. Gonorrhea is a global public health concern. In 2012 the World Health Organization (WHO) estimated over 78 million new urogenital cases per year in adults (15–49 years of age) worldwide (1, 2). The spread of gonorrhea is facilitated by the high prevalence of asymptomatic infections. Urogenital gonorrhea is asymptomatic in up to 10-15% of infected men and up to 50% of infected women. Pharyngeal and rectal infections, which have increased in prevalence in both sexes and are predominant among men who have sex with men, are primarily asymptomatic (3, 4). Untreated or inappropriately treated gonorrhea can result in serious consequences on reproductive and neonatal health. Women, in particular, are disproportionately affected, as gonococcal infection can ascend from the cervix to the uterus, Fallopian tubes, ovaries, and surrounding tissue, causing pelvic inflammatory disease. Long-term sequelae include ectopic pregnancy, chronic pelvic pain, and infertility. Furthermore, gonorrhea is strongly associated with an increased risk of both the acquisition and transmission of HIV (5).

Antimicrobial therapy is the only mainstay in the effective management and control of gonorrhea. However, *N. gonorrhoeae* exhibits an extraordinary capacity to develop antimicrobial resistance (AMR) through mutations and acquisition of AMR genes. The evolution of AMR in *N. gonorrhoeae* has overcome every therapeutic option since the “miracle drug” penicillin was introduced for gonorrhea treatment. Currently, a dual antimicrobial therapy (mainly ceftriaxone and azithromycin) is recommended for treatment of uncomplicated infections (6). Of grave concern over the past decade is the proliferation of resistance or decreased susceptibility to ceftriaxone worldwide. Azithromycin resistance has also emerged in most settings (7). The first failure of one of the recommended dual antimicrobial therapies against pharyngeal gonorrhea was reported in 2016 (8), and the first *N. gonorrhoeae* isolates with resistance to ceftriaxone combined with high-level resistance to azithromycin were identified in the United Kingdom (9, 10) and Australia (11, 12) in early 2018. In consideration of dwindling treatment options, scarce therapeutic alternatives, disease prevalence and morbidity, and lack of a vaccine(s), *N. gonorrhoeae* has been categorized by the WHO as a high priority pathogen globally and by the Centers for Disease Control and Prevention (CDC) as an urgent level threat in the USA (13).

Developing an effective gonococcal vaccine is essential because this is the only sustainable solution to quell the spread of gonococcal AMR and gonorrhea in general. The battle against penicillin-nonsusceptible *Streptococcus pneumoniae* exemplifies a successful vaccination strategy. Introduction of a pneumococcal conjugate vaccine in 2010 reduced the number of infections over 45% (14). Unfortunately, despite its public health importance, gonorrhea vaccine development remains in its infancy. Since 1970, only three small-scale vaccine trials using whole cell (15), pilin (16), and porin proteins (17) have been launched. All were unsuccessful in developing immunity against reinfection with gonorrhea. However, recent breakthroughs, including the development of small animal models for evaluating gonorrhea vaccines (18, 19), increased knowledge about *N. gonorrhoeae* immune evasion mechanisms (20–26), and the development of an effective vaccine for the closely related *N. meningitidis* serogroup B, which provided a low level of cross-protection against gonococcal infection (27), have reinvigorated the interest in gonococcal vaccine development (28).

Proteomic technology offers a powerful toolbox to enable vaccine antigen mining (28–32) and AMR proteome analysis (33–35), and to provide insights into host-pathogen interactions (36–39). Proteomic approaches have an advantage over genomics in drug and vaccine discovery endeavors by delivering information pertaining to protein abundance, post-translational modification(s), structure-function relationships, and protein-protein interactions (40–42). In addition, subcellular fractionation steps preceding proteomic applications reduce sample complexity, increase the likelihood of discovering low-abundance proteins, and aid in defining protein localization, all of which provide further insights into the proteins’ functions and interactomes (43, 44). For *N. gonorrhoeae*, proteomic approaches have begun to deliver proteinaceous vaccine candidates (29, 30, 39, 45) and to support elucidation of AMR patterns (46, 47). Current off-gel proteomics, such as isobaric tag labeling (isobaric tagging for absolute quantification, iTRAQ; and tandem mass tags, TMT) coupled with high-pressure liquid chromatography and mass spectrometry techniques (LC-MS/MS), demonstrate superb protein separation and identification and enable detection of proteins in the low femtomole to high attomole range with precision and reliability (29, 48, 49).

To address the need for discovery of additional gonorrhea vaccine and drug candidates and to enhance our understanding of AMR at the proteome level, herein we examined the 2016 WHO *N. gonorrhoeae* reference strains (50) and the FA6140 strain (51) using a global quantitative proteomic approach. The WHO panel consists of 14 *N. gonorrhoeae* reference strains strictly selected and validated internationally to represent the *N. gonorrhoeae* species. All known gonococcal phenotypic and genetic AMR determinants are included for use as quality control strains during phenotypic and genetic laboratory testing. Eight of the strains were initially included in the 2008 WHO reference strains [WHO F, G, K, L, M, N, O, and P; (52)] to which 6 novel strains (U, V, W, X, Y, and Z) were added to constitute the 2016 WHO reference strains (50). All WHO panel strains have been subjected to extensive phenotypic, genomic, and genetic analyses to establish diagnostic markers, molecular epidemiological characteristics, reference genomes, and AMR profiles (phenotypic and genetic) for all antimicrobials currently and previously used for gonorrhea treatment, in addition to novel antimicrobials considered for future interventions. This panel includes WHO X, the first extensively drug-resistant gonococcal strain identified with high-level resistance to ceftriaxone, as well as additional strains with different levels of resistance to ceftriaxone, azithromycin and any additional therapeutic antimicrobials. Complete genomes with detailed annotations are available for all panel strains, providing a fundamental resource for future molecular studies. Accordingly, the well-characterized 2016 WHO reference strains (50) are ideally suited to provide detailed descriptions of the global *N. gonorrhoeae* proteome, a greater understanding of gonococcal AMR at the proteome level, and a source for the identification of broadly conserved novel vaccine candidates. In addition to the WHO panel strains, we have included in our investigations *N. gonorrhoeae* FA6140, which is a penicillin-resistant, β-lactamase-negative isolate that was originally described after a local epidemic outbreak of 199 gonococcal cases in Durham, North Carolina, USA in 1983 (51). It serves as a model for gonococcal AMR studies and has facilitated the characterization of mutations in genes encoding the “multiple transferable resistance” repressor MtrR (53), ribosomal protein S10 (54), and penicillin-binding protein 2 (55) and their impact on AMR.

Our study is the first to investigate the global proteomic profiles of 15 *N. gonorrhoeae* reference strains using subcellular fractionation to separate cytoplasmic (C) and cell envelope (CE) associated proteomes, which were measured with tandem mass tags coupled to liquid chromatography and tandem mass spectrometry [TMT-LC-MS/MS; (56)], a highly reproducible and sensitive technique. These proteomic studies achieved our three major objectives. First, to enhance progress on gonorrhea vaccine development, novel vaccine candidates were identified, and the expression profiles of currently proposed antigens were established in diverse clinical isolates. Second, to broaden our understanding of AMR, proteomic signatures associated with AMR were defined by conducting a pairwise analysis of differentially expressed proteins to compare FA6140 and the 2016 WHO panel to WHO F, which possesses the largest genome and is susceptible to all relevant antimicrobials (50). Third, to facilitate the use of the 2016 WHO panel in various types of basic research and quality assurance, the complete reference proteomes of all tested strains were defined.

## EXPERIMENTAL PROCEDURES

### Bacterial strains and growth conditions

The 2016 WHO *N. gonorrhoeae* reference strains [n=14; (50, 52)] and the *N. gonorrhoeae* FA6140 strain (51) were used in this study. The AMR profiles of all isolates were described previously (50). Gonococcal strains were cultured from frozen stocks (−80°C) onto gonococcal base agar (GCB) medium (Difco) with Kellogg’s supplements I and II, diluted 1:100 and 1:1,000, respectively (57). After incubation at 37°C in a 5% CO_2_-enriched atmosphere for 18-20 h, nonpiliated and transparent colonies were subcultured onto GCB and incubated as described above. To initiate growth in liquid medium, nonpiliated colonies were collected from GCB and suspended to an OD_600_ of 0.1 in pre-warmed GCB liquid (GCBL) medium supplemented as described above with the addition of 0.042% sodium bicarbonate. Suspensions were incubated at 37°C with shaking at 220 rpm.

### Subcellular fractionation and TMT labeling

All 15 *N. gonorrhoeae* strains were simultaneously cultured in GCBL as described above. Cells were collected by centrifugation (20 min, 6,000 × *g*) when the Optical Density (OD_600_) of each culture reached 0.6 – 0.8, re-suspended in PBS and lysed by passage through a French Press. The cell debris was removed by centrifugation and the crude CE fraction was separated from the C proteins using a sodium carbonate extraction procedure and subsequent ultracentrifugation steps. The fraction enriched with CE proteins was reconstituted in PBS supplemented with 0.1% SDS (29, 30). Experiments were conducted in two biological replicates. Sample quality and the overall sub-proteome profiles were examined by SDS-PAGE coupled with Colloidal Coomassie staining (58, 59). The total protein amount in each fraction was assessed using a Protein Assay Kit (Bio Rad). Each CE and C fraction containing 100 μg of protein in 25 μL volume of triethylammonium bicarbonate buffer was reduced with tris(2-carboxyethyl)phosphine hydrochloride and the cysteines were alkylated using iodoacetamide. Proteins were digested using trypsin (Promega) at a 1:40 ratio. TMT reagents (ThermoFisher Scientific) were dissolved in acetonitrile (ACN) and used to label proteins in CE and C fractions as follows for the 10-plex experiment (ref 90111, Thermo Fisher Scientific): WHO F strain: TMT^10^-126, WHO K strain: TMT^10^-127C, WHO G strain: TMT^10^-127N, WHO M strain: TMT^10^-128C, WHO L strain: TMT^10^-128N, WHO O strain: TMT^10^-129C, WHO N strain: TMT^10^-129N, WHO U strain: TMT^10^-130C, WHO P strain: TMT^10^-130N, WHO V strain: TMT^10^-131; for the 6-plex experiment (ref 90402, Thermo Fisher Scientific): WHO F strain: TMT^6^-126, WHO W strain: TMT^6^-127, WHO X strain: TMT^6^-128, WHO Y strain: TMT^6^-129, WHO Z strain: TMT^6^-130, FA6140 strain: TMT^6^-131. Mixtures were incubated for 1 h at room temperature. The reaction was quenched by addition of 8 μL of 5% hydroxylamine. Samples were pooled, dried in a vacuum concentrator and stored at -80°C before separation by high pressure liquid chromatography (HPLC) and MS analysis.

### Sample fractionation and MS analysis

Samples were fractionated by strong cation exchange (SCX) with a Paradigm (Michrom Biosciences) HPLC with mobile phases of 5 mM potassium phosphate monobasic in 30% ACN/70% water (v/v) pH 2.7 (buffer A) and 5 mM potassium phosphate monobasic in 30% ACN/70% water (v/v) pH 2.7 with 500 mM potassium chloride (buffer B). The sample was brought up in buffer A (200 µL). The peptides were separated using a 2.1 mm x 100 mm Polysulfoethyl A column (PolyLC) over 60 min at a flow rate of 200 µL/min. The separation profile was as follows: hold 2% B for 5 min, 2% to 8% B in 0.1 min, 8% to 18% B in 14.9 min, 18% to 34% B in 12 min, 34% to 60% B in 18 min, 60% to 98% B in 0.1 min and hold for 10 min. Fractions were collected in 96-well microtiter plates at 1 min/fraction. Sixty fractions were pooled into 12 and dried using a speed vac. The samples were desalted using Oasis HLB 1cc cartridges. The cartridges were washed with 70% ACN/0.1% trifluoroacetic acid (TFA) and equilibrated with 0.1% TFA. Samples were loaded onto the cartridge in 0.1% TFA, washed with 0.1% TFA, and eluted in 1 mL 70% ACN/0.1% TFA. The samples were dried by vacuum centrifugation.

Desalted SCX fractions were analyzed by liquid chromatography electrospray ionization mass spectrometry (LC/ESI MS/MS) with a Thermo Scientific Easy-nLC II (Thermo Scientific) nano HPLC system coupled to a hybrid Orbitrap Elite ETD (Thermo Scientific) mass spectrometer. In-line de-salting was accomplished using a reversed-phase trap column (100 μm × 20 mm) packed with Magic C^18^AQ (5-μm 200Å resin; Michrom Bioresources) followed by peptide separations on a reversed-phase column (75 μm × 250 mm) packed with Magic C^18^AQ (5-μm 100Å resin; Michrom Bioresources) directly mounted on the electrospray ion source. A 90-minute gradient from 7% to 35% ACN in 0.1% formic acid at a flow rate of 400 nL/min was used for chromatographic separations. The heated capillary temperature was set to 300C and a spray voltage of 2750 V was applied to the electrospray tip. The Orbitrap Elite instrument was operated in the data-dependent mode, switching automatically between MS survey scans in the Orbitrap [automatic gain control (AGC) target value 1,000,000; resolution 120,000; and injection time 250 msec] with MS/MS spectra acquisition in the Orbitrap (AGC target value of 50,000; 15,000 resolution; and injection time 250 msec). The 15 most intense ions from the Fourier-transform full scan were selected for fragmentation in the higher-energy C-trap dissociation (HCD) cell by higher-energy collisional dissociation with a normalized collision energy of 40%. Selected ions were dynamically excluded for 30 sec with a list size of 500 and exclusion mass by mass width +/− 10ppm. HPLC and MS/MS analyses were performed in the Proteomic Facility at the Fred Hutchinson Cancer Center, Seattle.

### Proteomic data analysis

Data analysis was performed using Proteome Discoverer 1.4 (Thermo Scientific). The data were searched against WHO_F_CDS with the common Repository of Adventitious Proteins (cRAP, http://www.thegpm.org/crap/) fasta file. Trypsin was set as the enzyme with maximum missed cleavages set to 2. The precursor ion tolerance was set to 10 ppm and the fragment ion tolerance was set to 0.8 Da. Variable modifications included TMT 6Plex (+229.163 Da) on any N-Terminus, oxidation on methionine (+15.995 Da), carbamidomethyl on cysteine (+57.021 Da), and TMT 6Plex on lysine (+229.163 Da). Data were searched using Sequest HT. All search results were run through Percolator for scoring. Quantification was performed using the canned TMT 6plex or TMT 10plex methods through Proteome Discoverer with stringent criteria for protein identification including 1% False Discovery Rate (FDR), at least one unique peptide per protein, each identified peptide restricted to a single protein, and the score for every detected peptide of ≥1. Differential protein expression between CE and C fractions was determined by comparing the normalized total reporter ion intensities of groups using the WHO F protein expression profile as a reference.

### Bioinformatic Analysis

To detect potential homologous proteins, amino acid sequences of each identified *N. gonorrhoeae* vaccine candidate were downloaded and compared against the GenBank proteome database (https://www.ncbi.nlm.nih.gov/genbank/) using our in-house designed program based on the Reciprocal Best Blast Hit approach (60) using BLASTP with the following parameters: percentage identity ≥50%, and E-value ≤1.0 e-5.

Differential protein expression in four proteomics data sets (CE and C fractions in two biological replicates) was designated by fold changes ≥1.5 or ≤0.667 in reference to strain WHO F. Due to the variable nature of protein expression in *N. gonorrhoeae*, we took a conservative approach to designate protein expression and a protein was categorized as “up-regulated” or “down-regulated” solely when the fold change abundance was higher than 1.5 or lower than 0.667, respectively, to that of WHO F consistently in two biological experiments. A protein was designated as “ubiquitous” when its abundance was between 0.667-1.5-fold compared to WHO F in both experiments, or “variable” when its protein levels were not consistent between experiments.

A comprehensive assessment of predicted subcellular protein localization was accomplished by using the CELLO (61), PsortB 3.0.2 (62), SOSUI-GramN (63), SignalP 4.1 (64), LipoP 1.0 (65), and TMHMM 2.0 (http://www.cbs.dtu.dk/services/TMHMM/) prediction algorithms and a majority voting strategy. Furthermore, for proteins whose subcellular localization was not predicted using the aforementioned algorithms, we relied on the difference between their unique peptide counts in the CE and C fractions as follows:

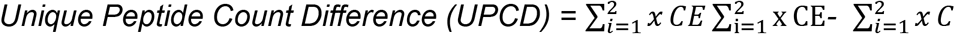

where “*i*” is the sequential number assigned for samples and “𝑥” is the total number of peptides detected in each fraction. Cytoplasmic proteins had more of their unique peptides detected in the C fraction (UPCD < 0), while membrane proteins had unique peptides enriched in the CE fraction (UPCD > 0). Proteins with UPCD=0 were excluded from analysis using this UPCD formula. Proteins were categorized as follows: outer membrane, periplasmic, inner membrane, C proteins, and proteins with unknown localization.

The phenotypic and proteotypic clusters of all strains were constructed using as variables both their AMR (50) and proteomic profiles obtained in this study. These clusters were designed based on the Hamming distance between tested strains, which counts how many elements differ between two vectors, and is equivalent to Manhattan distance on binary data. Average linkage was used to determine distances between clusters.

Graphs were generated with GraphPad Prism version 7 for Mac (GraphPad Software). The proteotypes of strains that belong to the same phenotypic cluster were compared, highlighting proteins that are significantly up-or down-regulated with respect to those proteins of WHO F.

### Data Availability

The raw mass spectrometry data have been deposited to the ProteomeXchange Consortium via the PRIDE (66) partner repository with the data set identifier PXD008412.

## RESULTS and DISCUSSION

### Study rationale

In our study design (Fig. 1), all 15 strains were cultured concurrently to mid-logarithmic growth, harvested, and subjected to subcellular fractionation to separate CE (outer membrane, periplasmic, inner membrane) and C proteins. We utilized TMT reagent technology for protein identification and quantitation as it provides a highly sensitive method for peptide labeling (56) and allows up to 10 biological samples to be analyzed in a single experiment (67). TMT-labeling, two-dimensional liquid chromatography fractionation, and subsequent MS/MS analyses were conducted on every 6-plex and 10-plex experiment pertaining to the CE and C fractions derived from each strain (Fig. 1). We selected WHO F as the reference strain for protein identification and quantitation because it has the largest genome (2,292,467 bp) and proteome (2,450 ORFs) among the 2016 WHO reference strains (50) and FA6140 (68), and it is susceptible to most antimicrobials currently or historically used for gonorrhea treatment.

**Figure 1.**
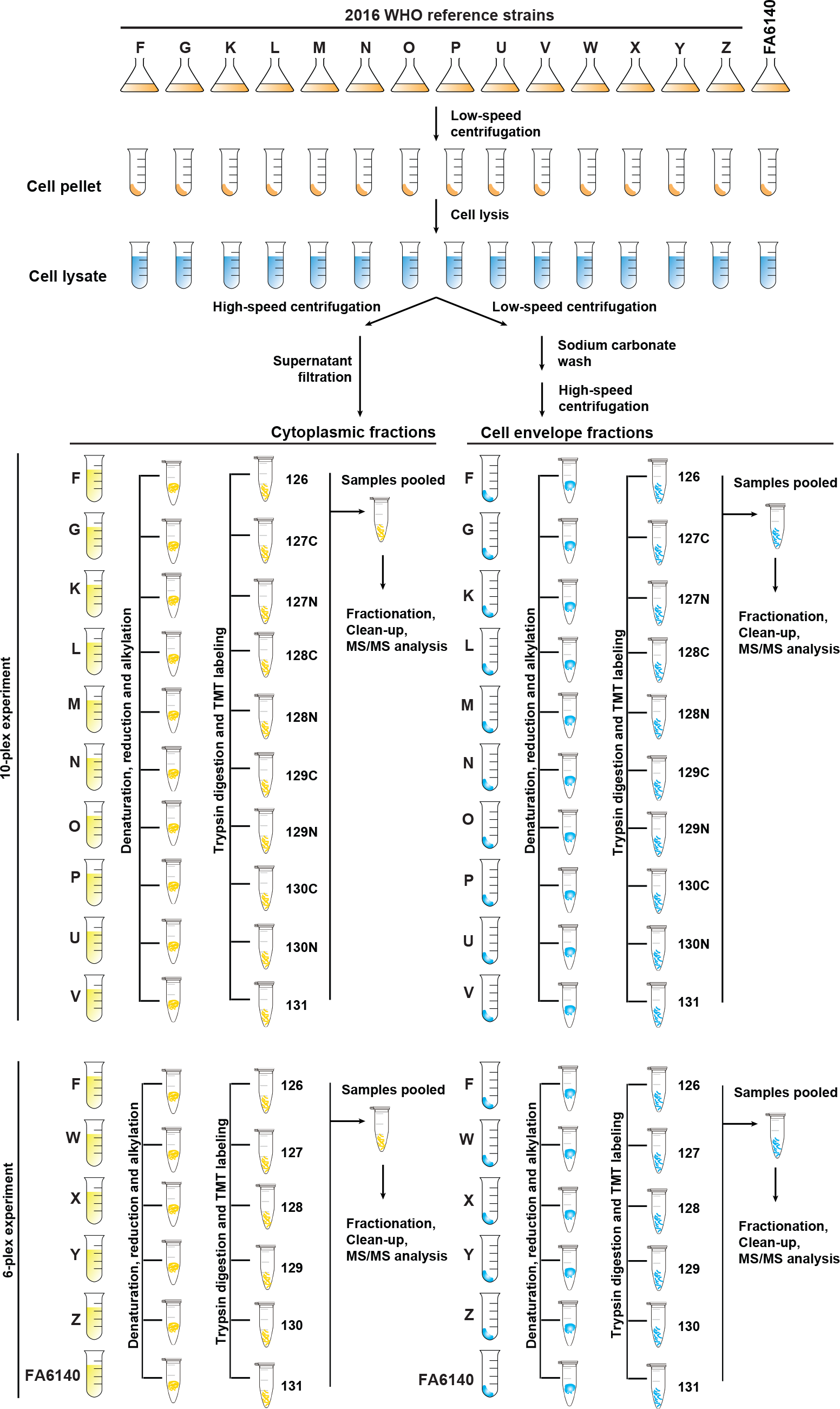
Experimental paradigm of quantitative proteomic profiling of the *N. gonorrhoeae* 2016 WHO reference strains and the FA6140 strain. All gonococci were cultured concurrently in liquid medium until reaching mid-logarithmic growth. Bacterial cells were harvested, lysed, and subjected to subcellular fractionation to separate the crude cell envelope (CE) and cytoplasmic (C) proteomes. CE proteins were enriched using a sodium carbonate wash and ultracentrifugation. The obtained CE and C protein samples (100 µg) were denatured, reduced, alkylated, trypsinized, and the peptides from each strain were labeled using 10-plex and 6-plex Tandem mass tag (TMT) reagents, as indicated. Finally, samples were pooled, fractionated by strong cation exchange, and analyzed by liquid chromatography electrospray ionization mass spectrometry. Experiments were performed in biological duplicates.

Sub-cellular fractionation experiments coupled with proteomics repeatedly show cytoplasmic proteins associated with the membranes, which are commonly regarded as “contaminating” or “moonlighting” proteins (29, 30, 69, 70). Therefore, to focus solely on the enriched proteins in individual subproteomes, we first eliminated C and CE proteins that were detected in the CE and C protein fractions, respectively, from further analyses. Complete lists of all identified proteins are in Supplemental Tables S1-S2. Subsequently, we performed two-armed proteomic data analyses: 1) for vaccine antigen mining, we focused on common proteins identified in all strains in the CE fraction with the overarching goal to discover omnipresent *N. gonorrhoeae* proteins; 2) to profile AMR signatures, we performed a pairwise comparison of individual strains to WHO F to enhance the discovery of strain-specific feature(s).

### Overview of cell envelope and cytoplasmic proteomes

The 10-plex biological replicate experiments identified a total of 1150 proteins in the CE fraction of all ten strains, of which 1010 were common in both sets (Fig. 2 A). In the two 6-plex experiments, 1194 proteins were identified; 975 were shared in all six isolates (Fig. 2 A). Taken together, the 10-plex and the 6-plex experiments resulted in identification of 1084 proteins in the CE fractions, of which 901 were common among all examined *N. gonorrhoeae* strains (Fig. 2 A). The proteome coverage per strain ranged from 41.22% (981 proteins) for WHO Y to 45.32% (1042 proteins) for WHO G (Supplemental Table S3).

**Figure 2.**
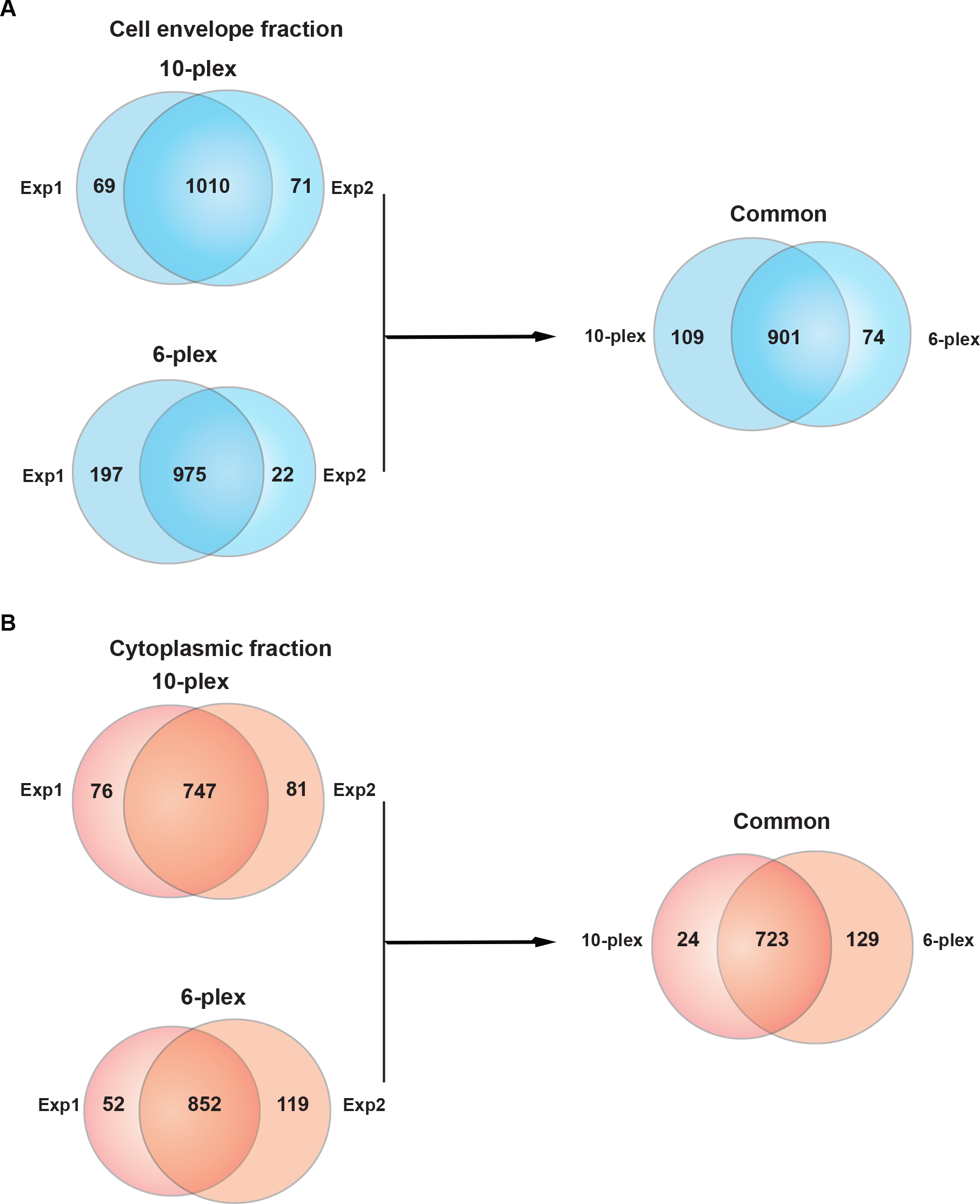
Venn diagrams illustrating the distribution of proteins identified in cell envelope and cytoplasmic fractions in two independent proteomic experiments. **(A)** Cell envelope proteomes derived from the 2016 WHO reference strains and FA6140 were analyzed in 10-plex and 6-plex experiments performed in biological duplicates. A total of 1079 and 1081 proteins was identified in Experiments 1 and 2, respectively, and 1010 common proteins were found in both 10-plex experiments. The 6-plex TMT labeling revealed 975 common proteins as well as 197 and 22 unique proteins in Experiments 1 and 2, respectively. Further analyses were applied to 901 proteins mutually identified in both 10-plex and 6-plex experiments. **(B)** The proteomic profiling of cytoplasmic fractions yielded 904 proteins shared among all 10 strains, of which 747 were common in both experiments. The 6-plex TMT identified 904 and 971 proteins in Experiments 1 and 2, respectively; of Proteomic mining of gonorrhea antigens and AMR which 852 were common between replicates. In further analyses solely the 723 proteins shared between both experiments were included. Exp 1 - experiment 1; Exp 2 - experiment 2.

Proteomics of the C fraction in the 10-plex set conducted in biological replicates yielded 904 proteins that were shared among all 10 strains, of which 747 were common in both experiments (Fig. 2 B). The two 6-plex experiments identified 1023 shared proteins, with 852 common among the two replicates (Fig. 2 B). Cumulatively, C fraction profiling resulted in identification of 876 proteins with 723 common in all 15 *N. gonorrhoeae* strains (Fig. 2 B). Proteome coverage ranged from 31.37% (746 proteins) in WHO U to 38.43% (852 proteins) in FA6140 (Supplemental Table S3).

Subsequently, we allocated common proteins that were identified in all 15 *N. gonorrhoeae* strains to outer membrane, inner membrane, periplasm, cytoplasm, or unknown localization categories based on PSORTb 3.0.2 (62), SOSUIGramN (63), and CELLO (61) predictions and the majority-voting strategy. We used these software packages to take advantage of their different algorithms and statistical approaches for the prediction of protein subcellular localization. As expected from our subcellular fractionation approach (49, 69, 71), the CE fraction was enriched in membrane proteins in comparison to the C sample, with outer membrane (26 vs. 8), periplasmic (51 vs. 38), and inner membrane proteins (145 vs. 6) that were also identified with considerably higher peptide counts (Fig. 3 A-C, and Supplemental Tables S1-S2, and S4-S5). The C preparations yielded 592 cytoplasmic proteins that were identified with greater peptide counts in comparison to the cytoplasmic proteins associated with the CE fraction (Fig. 3 D, Supplemental Tables S5-S6). Furthermore, to increase the discovery of potential vaccine candidates, we searched the 149 proteins of unknown localization identified in the CE fraction (Fig. 3 D) for the presence of signal peptides and transmembrane motifs using SignalP 4.1 (64), LipoP 1.0 (65), and TMHMM 2.0 (http://www.cbs.dtu.dk/services/TMHMM/). The results of all software programs and the majority votes strategy revealed six additional CE proteins, four of which were found in the majority of examined strains (Supplemental Table S6). In addition, literature searches for experimental evidence of protein surface exposure allowed assignment of BamE [NGO1780; (72)], SliC [NGO1063; (73)], MetQ [NGO2139; (30, 74)], Ng-MIP [NGO1225; (30, 75)], and BamG [NGO1985; (76)] to the cell surface.

**Figure 3.**
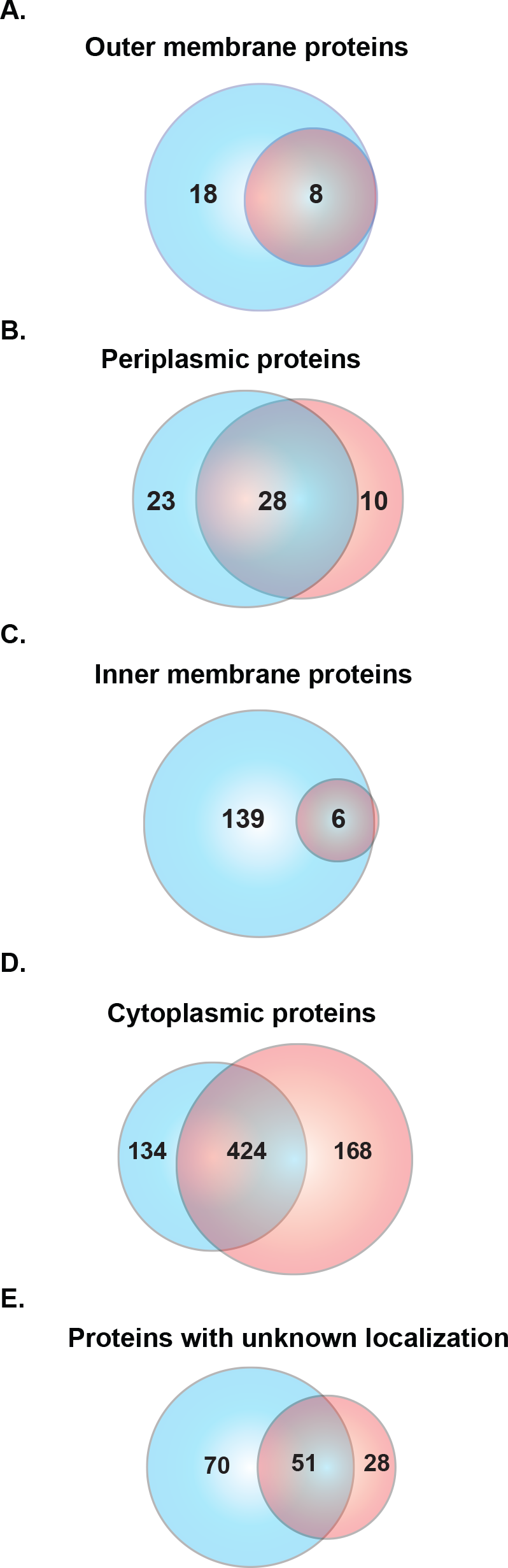
Subcellular localization of proteins identified in cell envelope and cytoplasmic subproteomes. Proteins identified in the cell envelope (blue circle) and cytoplasmic (red circle) fractions were subjected to comprehensive assessments of subcellular localization using different prediction algorithms and were allocated into the outer membrane **(A)**, periplasm **(B)**, inner membrane **(C)**, cytoplasm **(D)**, or unknown localization **(E)**.

### Expression patterns of common identified proteins in comparison to WHO F

Proteins were categorized as ubiquitous, up-or down-regulated, or variable based on their abundance in relation to the corresponding protein in WHO F in biological duplicate experiments. We investigated expression patterns of detected proteins in both sub-proteome fractions that were shared among all strains (Figs. 4–5, Supplemental Tables S4-5). Annotated cell envelope proteins were predominantly ubiquitous in the CE fraction. The proportion of ubiquitous CE outer membrane proteins (*n*=26) ranged from 73% (*n*=19 in WHO N) to 46.15% (*n*=12 in WHO L; Fig. 4 A). Ubiquitous periplasmic proteins ranged from 76.47% (*n*=39 in WHO N) to 35.2% (*n*=18 in WHO U) of the total number of proteins annotated to localize to the periplasm (*n*=51; Fig. 4 B). Finally, between 83.4% (*n*=121 in WHO M) and 44.1% (*n*=64 in WHO U) of inner membrane proteins (*n*=145) were ubiquitously expressed (Fig. 4 C). Up-regulated outer membrane (0 – 19.23%), periplasmic (0 – 8%) and inner membrane (0 – 4.8%) proteins made up the smallest proportion of proteins in the CE fraction (Supplemental Table S4), whereas down-regulated outer membrane (3.85 – 26.9%), periplasmic (0 – 23.53%) and inner membrane (1.38 – 24.8%) proteins were moderately prevalent (Supplemental Figure S4). Further analysis of the CE fraction detected 121 common proteins with unknown localization (Fig. 3 E, Supplemental Table S4). Ubiquitous expression was the dominant pattern for these proteins in WHO G, K, M, N, V, W, and X and ranged from 42.97% (*n*=52, WHO V) to 66.94% (*n*=81, WHO W). Variable expression of proteins with unknown localization dominated in WHO L, O, P, Y, Z, and FA6140. WHO U had the highest number of down-regulated proteins (39.67%, *n*=48) with respect to those in WHO F (Fig. 4 D).

**Figure 4.**
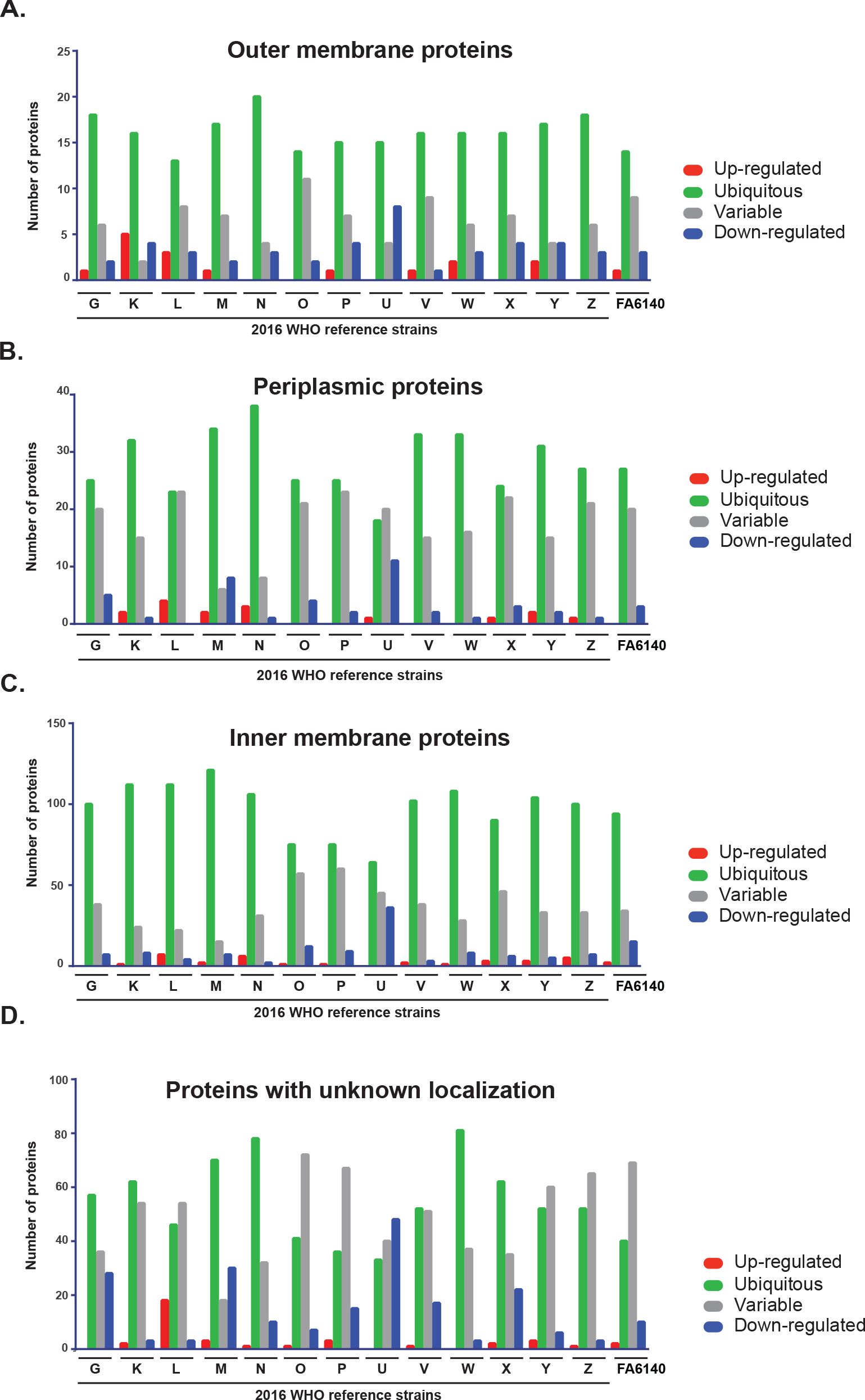
Expression patterns of common proteins identified in the cell envelope fraction. Outer membrane **(A)**, periplasmic **(B)**, inner membrane **(C)**, or proteins with unknown localization **(D)** are shown. Expression of each protein in each gonococcal strain was compared to the protein level in the reference WHO F isolate. Protein expression is categorized as ubiquitous (green bars); up-regulated (red bar); down-regulated (blue bar); and variable (grey bar).

In contrast to the CE expression pattern, we observed a striking increase in the number of variably expressed cytoplasmic proteins in the C fraction of analyzed strains in comparison to WHO F (Fig. 5 A, Supplemental Table S5). The percentage of variable proteins ranged from 32.9% to 82.6% for WHO G and WHO Y, respectively (Supplemental Table S5). Ubiquitous proteins were the next most common category and oscillated from 15.5% in WHO Y to 56.4% in WHO G. The third group contained up-regulated proteins (0 – 21.28%), and down-regulated proteins ranged from 0.5 – 3.88% (Supplemental Table S5). For proteins with no assigned localization, variable expression was the most prevalent pattern in WHO K, L, O, P, U, W, X, Y, Z, and FA6140 (Fig. 5 B), ranging from 79.75% (*n*= 63, WHO Y) to 46.83% (*n*= 37, WHO L). Ubiquitous proteins were the dominating group in WHO G, M, N, and V (Fig. 5 B). Finally, up-and down-regulated proteins constituted up to 11.39% and 7.59%, respectively, of the total proteins with unknown localization in the C fraction.

**Figure 5.**
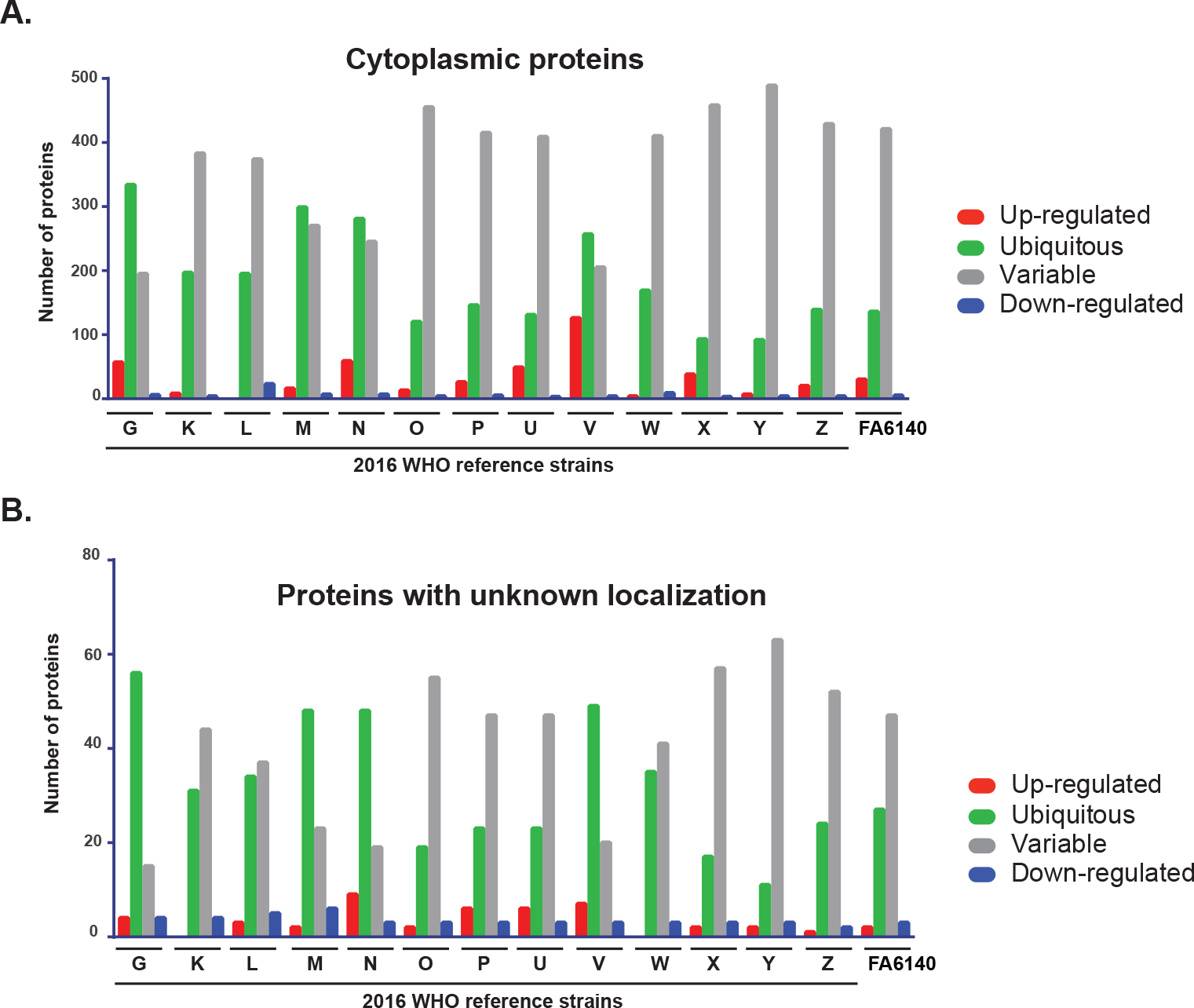
Expression patterns of common proteins identified in the cytoplasmic proteome. Cytoplasmic **(A)** and proteins with unknown localization **(B)** are shown. Protein levels in individual gonococcal strains were compared to the protein level in the reference Proteomic mining of gonorrhea antigens and AMR WHO F isolate. Protein expression is categorized as ubiquitous (green bars); up-regulated (red bar); down-regulated (blue bar); and variable (grey bar).

Together, the first quantitative proteomic profiling of the 15 *N. gonorrho*eae strains demonstrated distinct differences in their proteomes and showed that a pattern of ubiquitous protein expression was prevalent in the CE fraction, whereas variably expressed proteins were the dominant group in the C subproteome.

### Antigen mining decision tree

To identify novel gonorrhea antigens and to gain information about expression of previously identified vaccine candidates (49, 71, 77, 78), we designed an antigen mining decision tree (Fig. 6). We included in this process 25 outer membrane proteins (all outer membrane proteins identified except RmpM) and 121 proteins of unknown localization identified in CE proteomic profiling (Fig. 6 and Supplemental Table S1). The latter group of proteins was subjected to signal sequence and transmembrane motif analyses to increase the coverage of potential vaccine candidates. Together, these investigations yielded nine novel antigens including NGO0282, NGO0425, NGO0439, NGO0778, NGO1251, NGO1688, NGO1889, NGO1911a, and NGO2105 in addition to previously discovered proteomics-derived antigens [(29, 30); Table 1] and vaccine candidates identified by other means (Table 2).

**TABLE 1.**
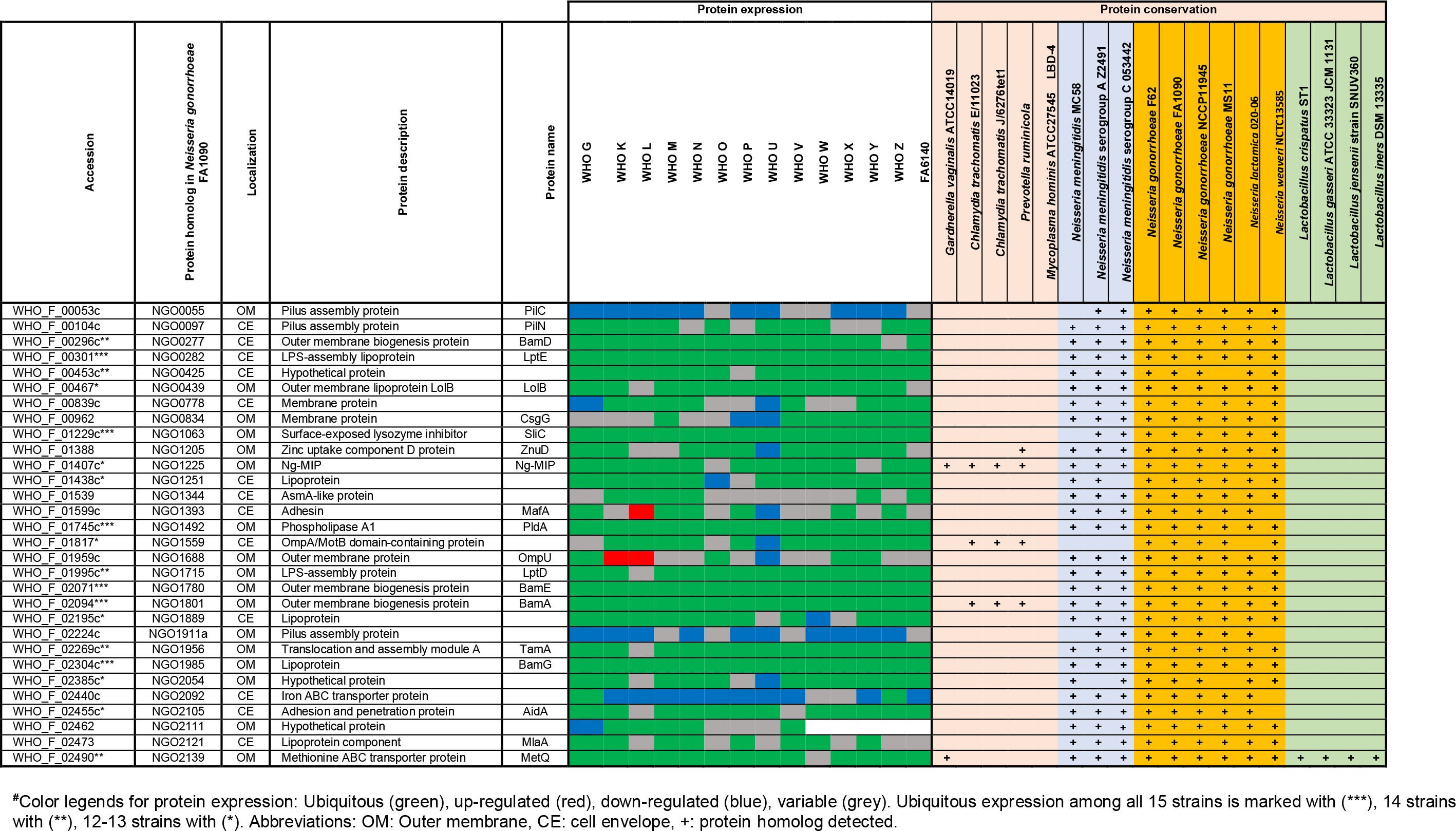
Expression and homologs of proteomics-derived *Neisseria gonorrhoeae* vaccine candidates#.

**TABLE 2.**
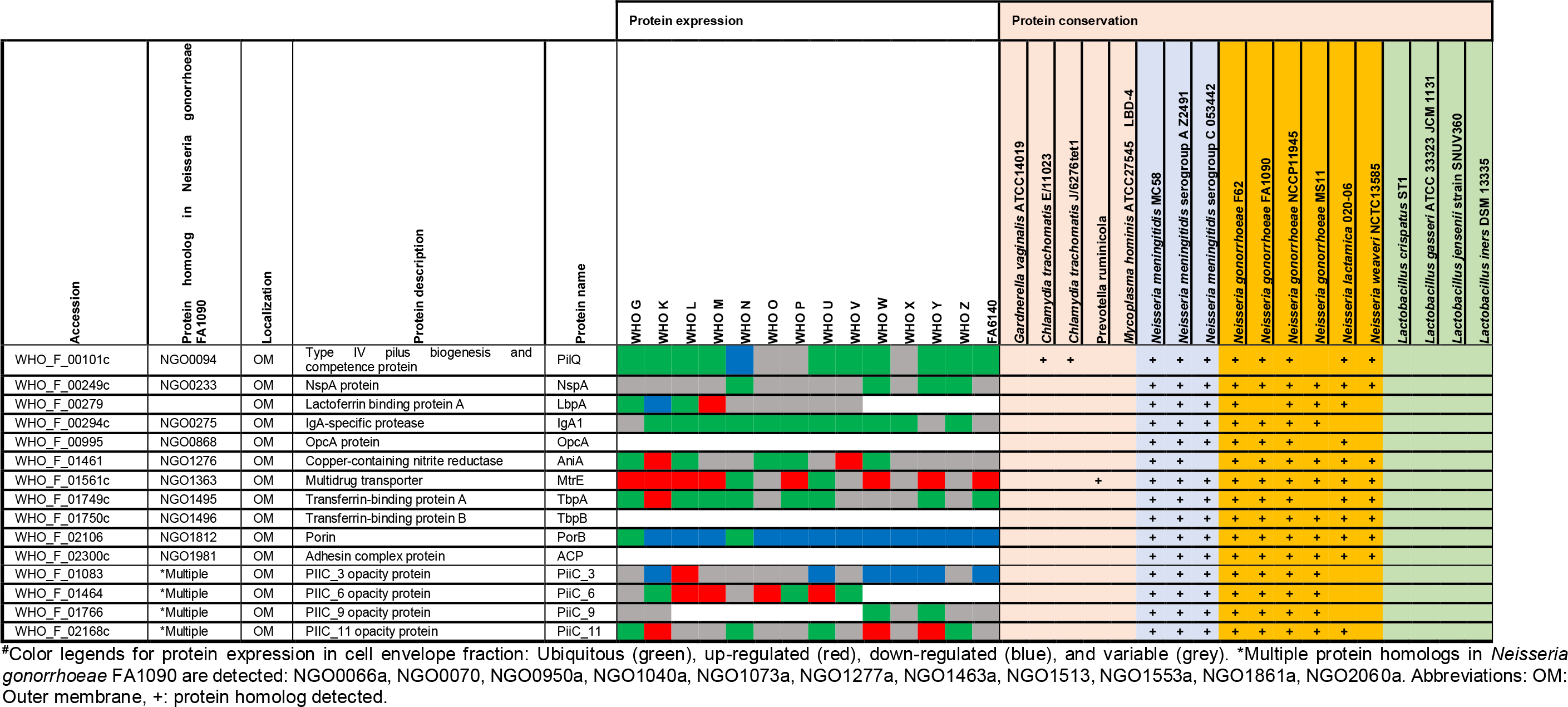
Expression and homologs of *Neisseria gonorrhoeae* vaccine candidates identified by other means#.

**Figure 6.**
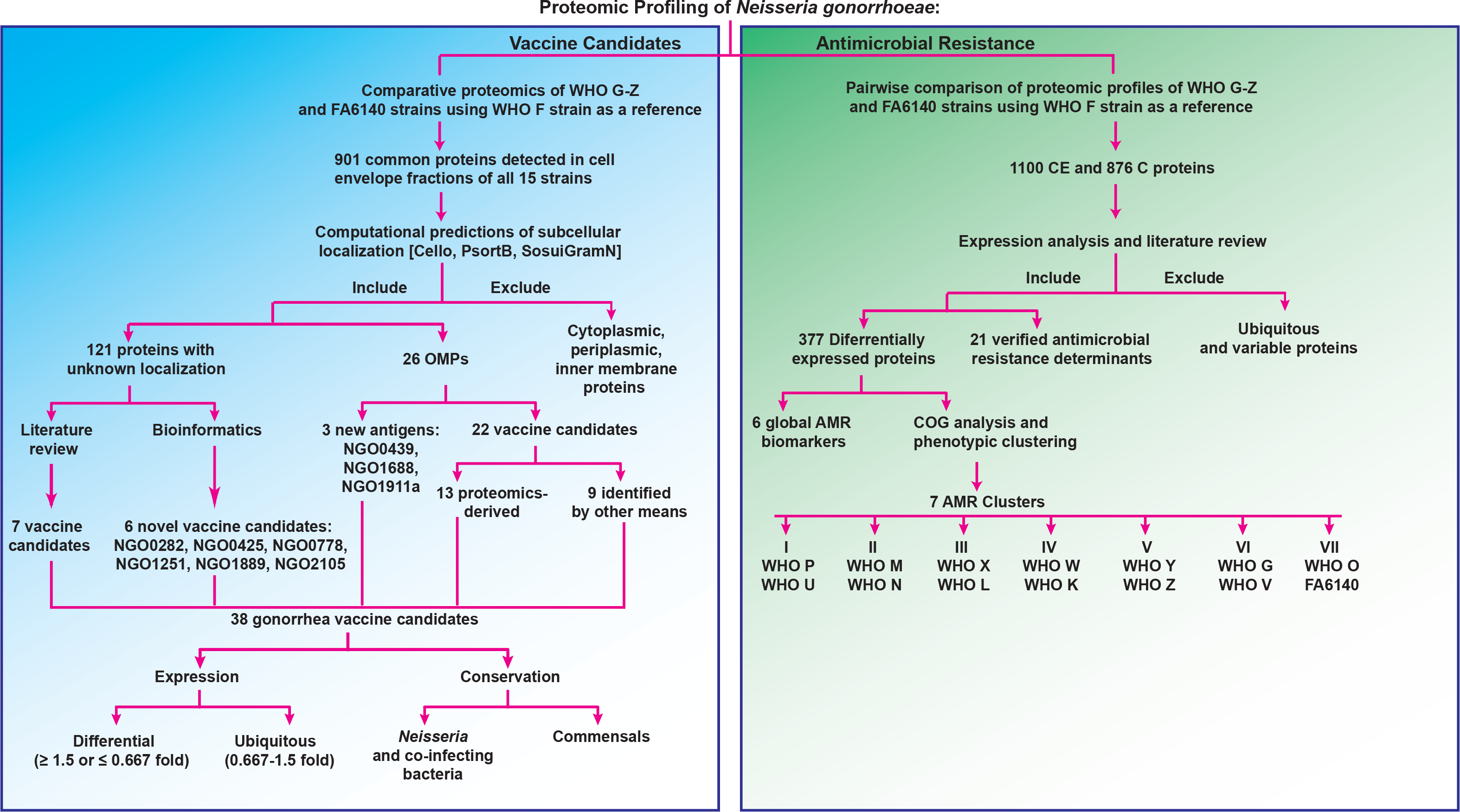
Decision tree designed for proteomic mining of *Neisseria gonorrhoeae* vaccine candidates and antibiotic resistance markers. Detailed description is provided in the text.

Further bioinformatics and literature searches were performed to gain insights into the new proteomics-derived vaccine candidates. The putative lipoprotein NGO0282 is a homolog of the outer membrane localized LptE, which is a component of the trans-envelope LptA-G machinery involved in the transport of lipopolysaccharide/lipooligosaccharide (LPS/LOS) molecules to the *E. coli* and *N. meningitidis* outer membrane, respectively. LptE’s chaperone-like role in LptD biogenesis is conserved in both bacteria but LptE works in concert with LptD to translocate LPS to the cell surface only in *E. coli* (79, 80). LptD is essential for *E. coli* and *N. gonorrhoeae* viability but is dispensable for *N. meningitidis* (81–83). Therefore, the function of *N. gonorrhoeae* LptE in both LOS transport and LptD biogenesis needs to be elucidated. NGO0425 contains a tetratricopeptide repeat–like domain and a transmembrane helix. Together with its *E. coli* homolog, YfgM, NGO0425 belongs to the UPF0070 family. In *E. coli*, YfgM was proposed to act within the β-barrel trafficking chaperone network and its depletion in Δ*skp* and Δ*surA* knockout backgrounds contributed to further alterations in outer membrane integrity (84). The vaccine candidate protein NGO0439 is homologous to the *E. coli* outer membrane protein LolB, which is involved in lipoprotein trafficking to the outer membrane (85, 86). We consider *N. gonorrhoeae* LolB to be a vaccine candidate antigen because its surface localization should be experimentally verified. Differences in the localization of homologous proteins exist. For instance, fHbp is a surface-displayed lipoprotein in *N. meningitidis* but not in *N. gonorrhoeae* (87), while BamE is on the surface of gonococci but faces the periplasmic side of the outer membrane in *E. coli* (72). Protein NGO1688, annotated as OmpU, is a putative iron uptake outer membrane protein that is positively regulated by the oxygen-sensing transcription factor, FNR (88). NGO1911a is a predicted pilus assembly protein that is associated with the adhesin PilY (89). Finally, NGO0778, NGO1251 (a putative lipoprotein), and NGO1889 are hypothetical proteins. NGO1889 belongs to the LprI family (PFO7007) that comprises bacterial proteins of ~120 amino acids in length that contain four conserved cysteine residues. LprI from *Mycobacterium tuberculosis* acts as a lysozyme inhibitor (90), providing the exciting possibility that *N. gonorrhoeae* LprI contributes to residual lysozyme resistance observed in gonococci deprived of surface-exposed lysozyme inhibitors SliC and ACP (73, 91). Lastly, NGO2105 contains peptidase S6 (residues 43-310) and autotransporter (residues 1215-1468) domains potentially involved in proteolytic activity and auto-translocation, respectively, suggesting that this is a newly identified autotransporter protein in *N. gonorrhoeae*. In support of this notion, the NGO2105 locus, also known as adhesion and penetration protein or “NEIS1959 (iga2)” in the PubMLST database, encodes IgA2 protease (AidA) and has homologs in other *Neisseria* (Table 1) as well as *Haemophilus influenzae* (92).

Together, our investigations yielded nine novel gonorrhea vaccine candidates, including proteins with implications in pathogenesis such as IgA2 (AidA) and LprI, and provided valuable information regarding the expression patterns of previously selected vaccine candidates.

### Expression and homologs of gonorrhea vaccine candidates

We first evaluated expression profiles of extensively studied gonorrhea vaccine candidates including MtrE (93–95), PorB (96, 97), PilQ (98), TbpA (99, 100), Opa (101, 102), and AniA (19, 103–105). MtrE and PorB were up-and down-regulated, respectively, in 12 isolates (Table 2). Compared to WHO F, PorB was present at similar levels only in WHO G and N. PilQ (98) was ubiquitously expressed in 10 strains, whereas expression of Opa proteins was widely variable, as expected (106, 107). The TbpA level was similar in 8 strains; however, we did not detect TbpB (108). Nor did we detect ACP (109, 110) or OpcA (111, 112) under the standard growth conditions used in our studies, which suggested that they might be specifically regulated. AniA was present at different levels in 7 strains, ubiquitous in five, and up-regulated in two isolates. Immunoblotting experiments with anti-AniA antisera corroborated these findings (105). The cellular pool of NspA (113) varied in ten isolates, while lactoferrin binding protein LbpA (114) was variable in five strains and was ubiquitous in WHO L and G (Table 2).

Strikingly, most of the proteome-derived vaccine candidates showed ubiquitous expression among numerous strains (Table 1). In particular, SliC, PldA, BamE, BamA, and BamG were ubiquitous in all 15 isolates. Similar results for these proteins were obtained by iTRAQ-MS/MS applied to the proteomic profiling of cell envelopes and outer membrane vesicles (OMVs) isolated from four different strains of *N. gonorrhoeae* (29). Further, LolB, Ng-MIP, NGO1559, and NGO2054 were unvaryingly expressed in at least 12 isolates. Among the novel vaccine candidates identified in our study, LptE, LolB, IgA2, and NGO1251 were found ubiquitous in at least 13 strains. In addition, LprI and NGO0778 were similarly expressed in 12 and 9 isolates, respectively. In support of our proteomics data, immunoblotting analyses demonstrated similar cellular levels of BamA, MetQ, TamA, LptD, NGO2054 (30), BamE-D (72), SliC (73), and BamG (76) in whole cell lysates of the 2016 WHO strains as well as geographically and temporally diverse clinical isolates of *N. gonorrhoeae* from Baltimore (*n=*5) and Seattle (*n=*13). Our previous studies showed that PorB, PilQ, BamA-D, SliC, MafA, PldA, MetQ, IgA1 protease, and LptD are cargo proteins present at similar levels in naturally released gonococcal OMVs (29, 72, 73), which further highlights their potential as vaccine antigens considering the success of *N. meningitidis* OMV-based vaccines (26, 115).

Finally, we examined the presence of homologs of the gonorrhea vaccine candidates among non-gonococcal *Neisseria* species, other commensal bacteria (116, 117), and co-infecting microbes (118–121) that inhabit the same ecological niche as *N. gonorrhoeae*. Antigens conserved between these pathogens and preferably not in commensals have the potential to eradicate several sexually transmitted infections, if formulated into a protective vaccine(s). Our comparative analyses showed that all of the proteomics-based antigens have homologous proteins in the majority of investigated *N. meningitidis* strains, and none are present in *M. hominis* (Table 1). In addition, Ng-MIP-like proteins exist in *C. trachomatis*, *G. vaginalis*, and *P. ruminicola*; BamA and NGO1559 homologs were found in *C. trachomatis* and *P. ruminicola*. MetQ, a methionine transporter (74), was the only proteomics-derived vaccine candidate with homologs across all examined bacteria with the exception of *C. trachomatis* and *P. ruminicola*. Further, we detected protein homologs of PilQ in two *C. trachomatis* strains and MtrE and ZnuD in *P. ruminicola*; these three proteins were absent in commensal species.

Together, our investigations provide pioneering information into newly identified and existing gonorrhea vaccine candidates. We have established each candidate’s expression pattern in diverse *N. gonorrhoeae* isolates and identified homologs among other pathogenic and/or commensal bacteria that share the same ecological niche. Stable expression in the WHO gonococcal panel coupled to presence in *N. meningitidis* and co-infecting agents – but rarely in urogenital commensals – further highlights the importance of including these antigens in gonorrhea vaccine(s).

### Proteomics profiling of N. gonorrhoeae antimicrobial resistance

Various genome-based AMR determinants have been deciphered in the gonococcus over the past decades (51, 122–127). However, many AMR determinants remain to be identified and characterized, e.g. the chromosomally-encoded penicillin and cephalosporin resistance determinant “factor X” (128–130) and the AMR mechanisms that contribute to a large proportion of azithromycin resistance (131). The uncertainty behind these AMR determinants illustrates the need for alternative approaches to enhance our understanding of gonococcal AMR complexity. At the proteomic level, only two studies have attempted to address this challenge, both of which used 2D-SDS PAGE exclusively (47, 132). Therefore, we focused on identifying proteomic AMR signatures that exist in the absence of antimicrobial pressure during standard *in vitro* growth conditions by performing a pairwise comparison of all identified proteins in each individual strain to the WHO F reference strain (Supplemental Tables S1-S2). As expected, we identified different numbers of proteins in the CE and C fractions in each comparison set due to differences in the number of open reading frames (ORFs) between the gonococcal strains (Supplemental Table S3). Similarly to our previous analysis, we excluded typical cytoplasmic proteins from the CE subproteome and cell envelope proteins from the C fraction. We solely focused on proteins with significantly different expression in the examined strains compared to the fully antimicrobial-susceptible strain WHO F with the rationale that these proteins may provide clues about the proteomic basis of AMR. For instance, we identified MtrE as up-regulated in many strains with increased resistance to numerous antimicrobials even in the absence of antimicrobial exposure, which represents an up-regulation of the multidrug MtrCDE efflux pump and possibly additional efflux pumps for which MtrE acts as the outer membrane channel (29, 94, 133–135). Overall, we identified 162 (including 21 known AMR determinants) and 95 proteins with known and unpredicted subcellular locations, respectively (Figure 6). Peptide counting performed for the latter group of proteins yielded 55 and 36 proteins that are likely localized to the CE and C, respectively, and four proteins with ambiguous localization. Next, we separated proteins that have been previously verified as *N. gonorrhoeae* AMR determinants (Table 3) from new potential proteomic AMR signatures (Tables 4–5).

**TABLE 3.**
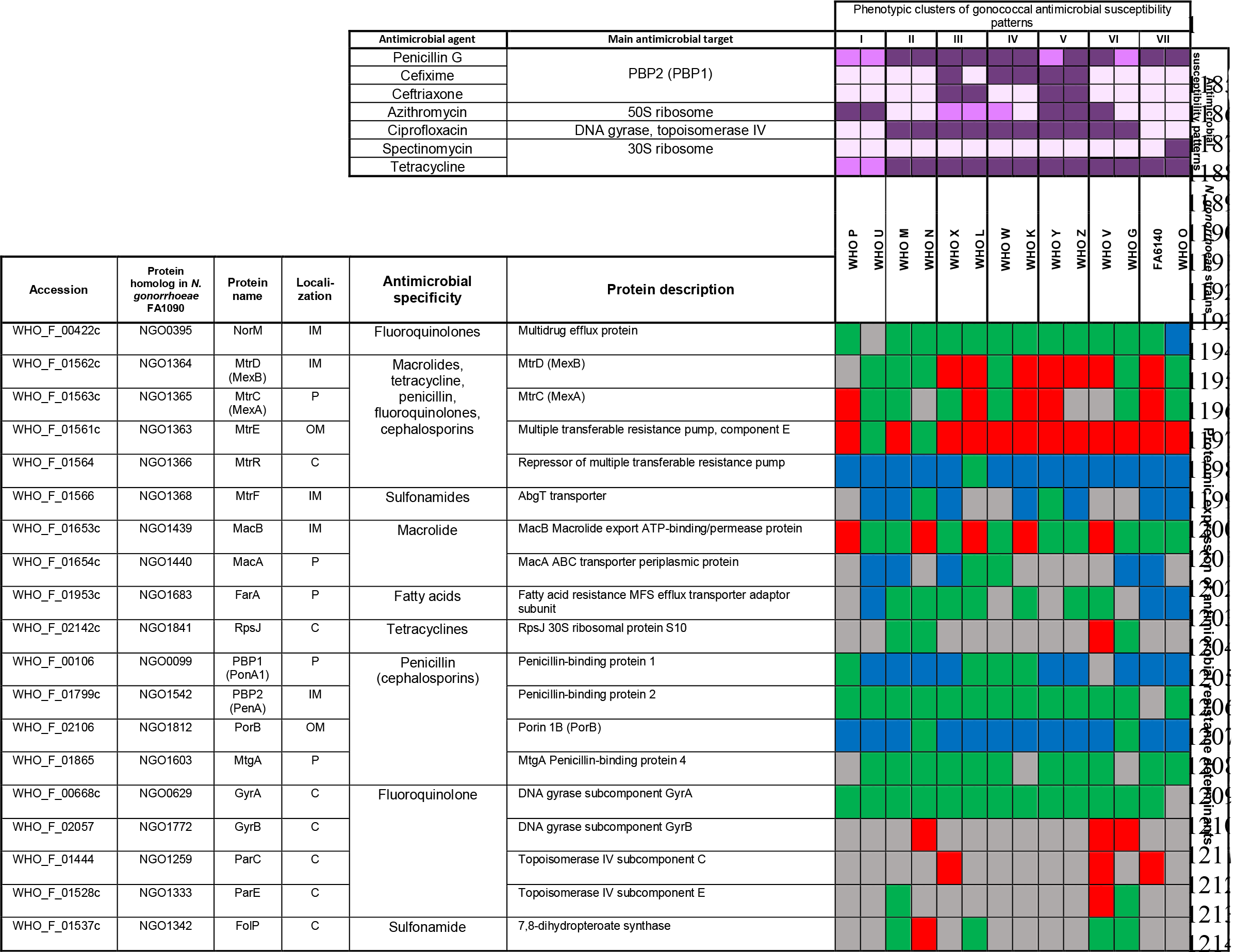
Proteomic signature of previously verified gonococcal antimicrobial resistance determinants#.

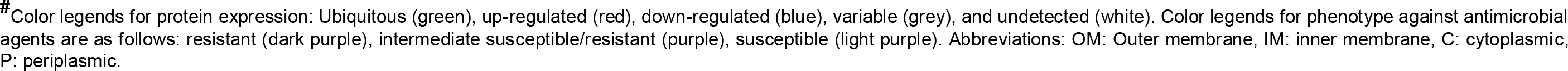

**TABLE 4.**
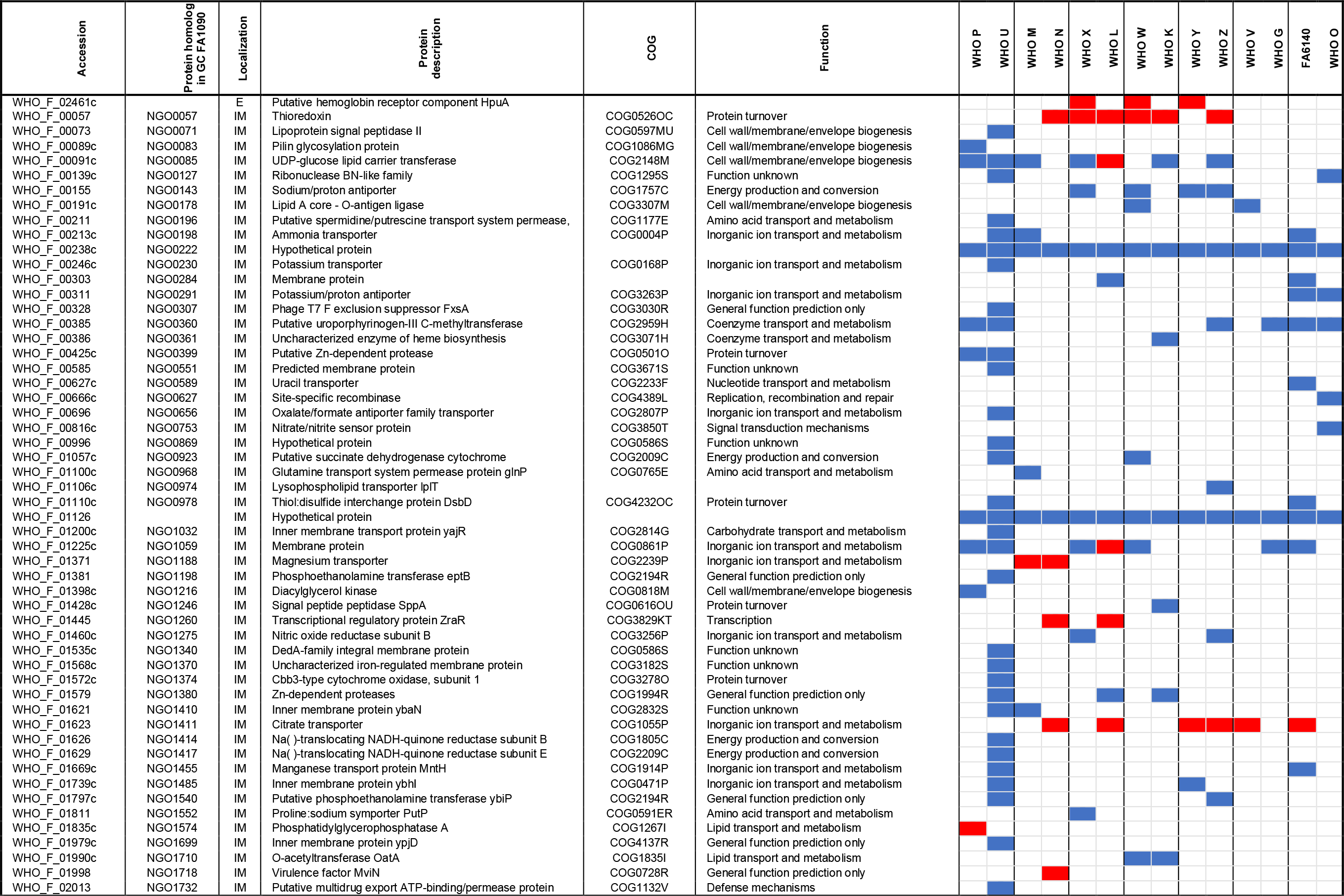
New potential proteomic-derived antimicrobial resistance signatures with defined subcellular localizations#.

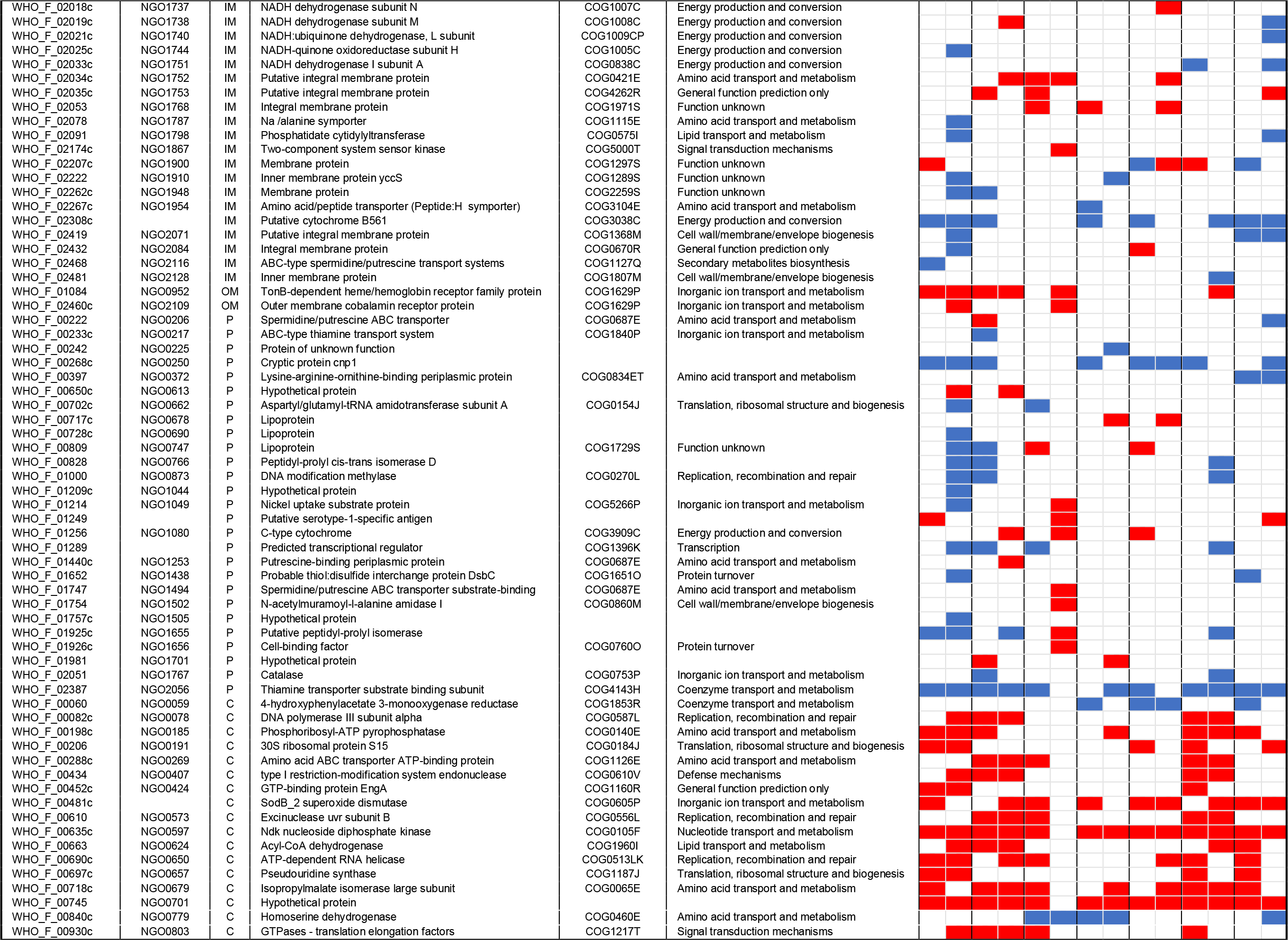

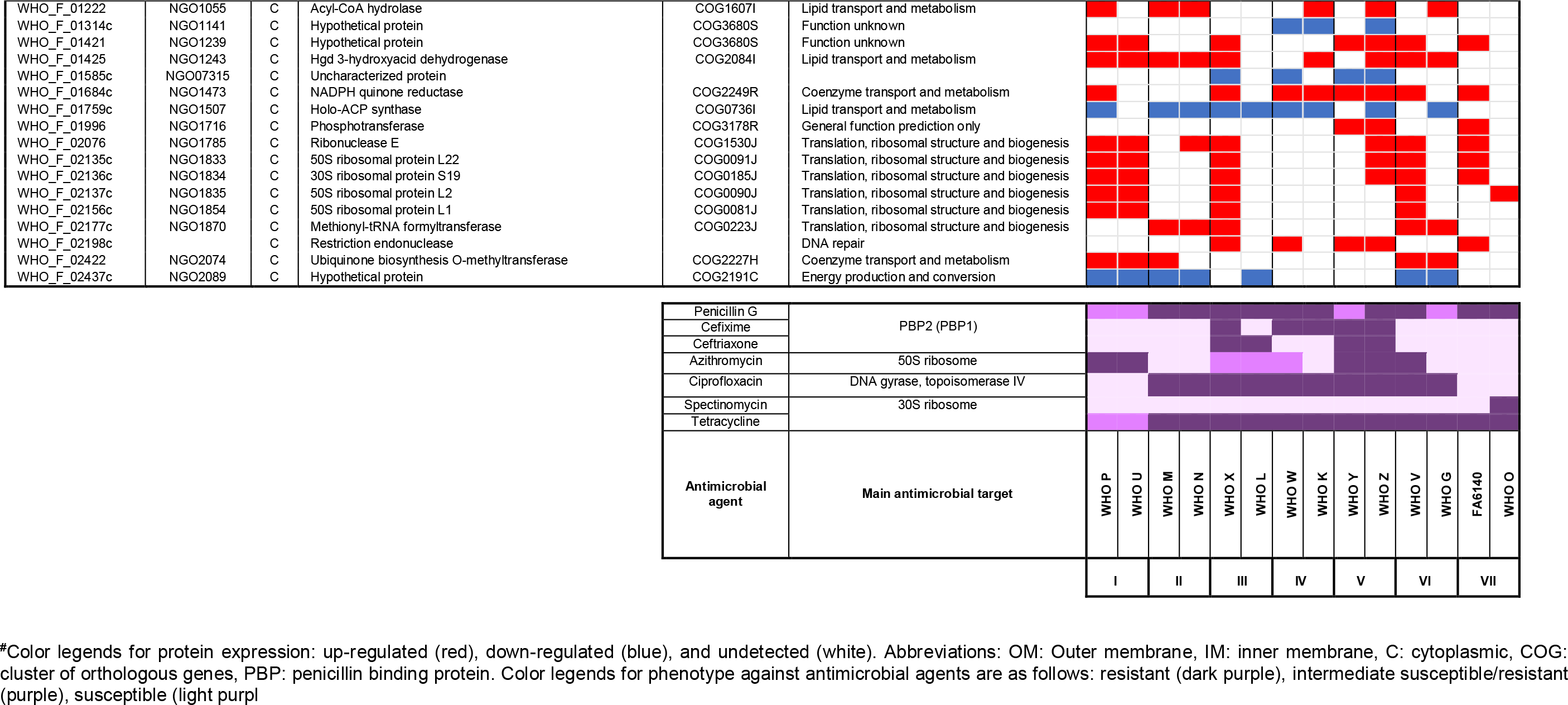

**TABLE 5.**
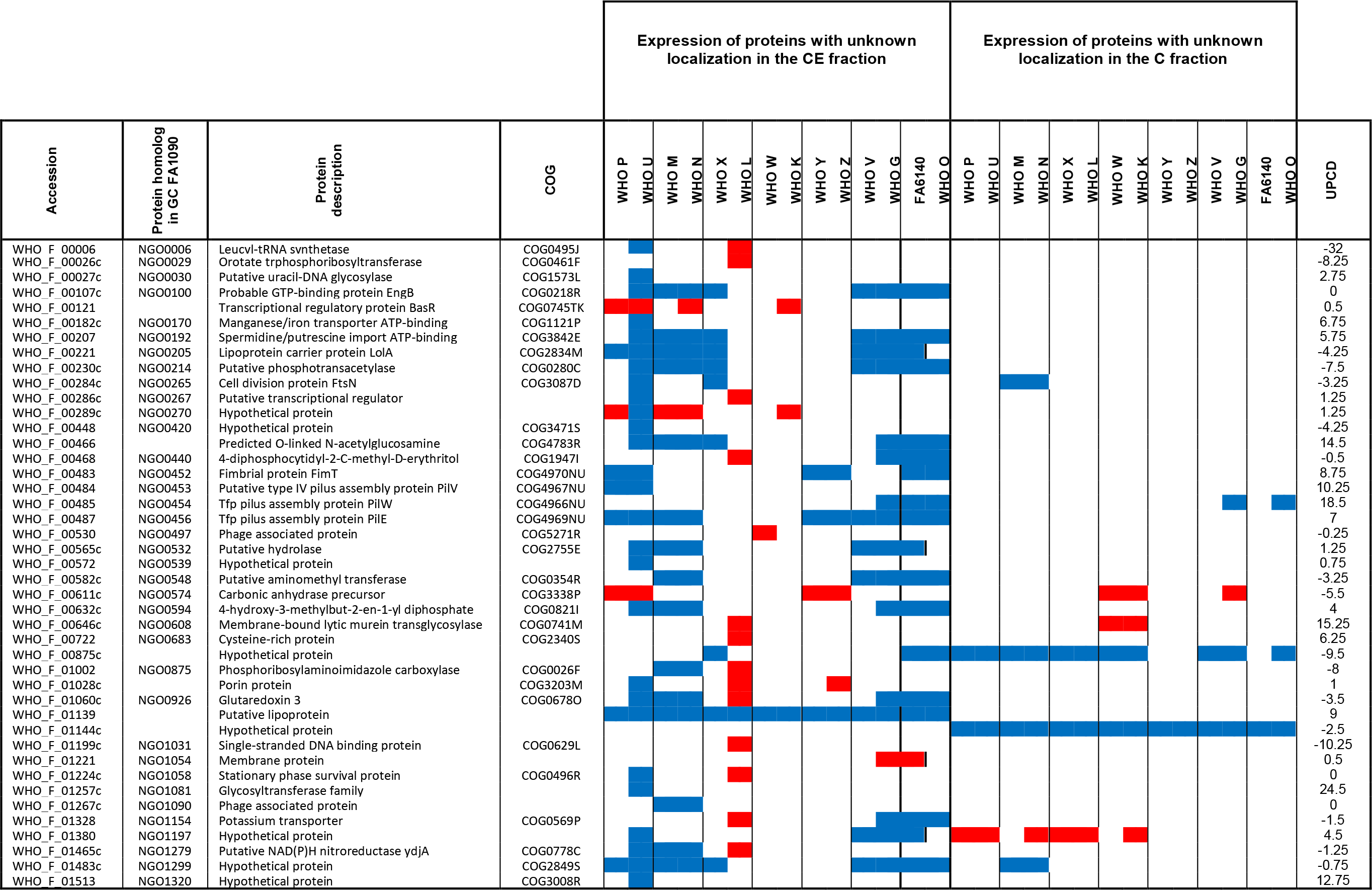
New potential proteomic-derived antimicrobial resistance signatures with undefined localization#.

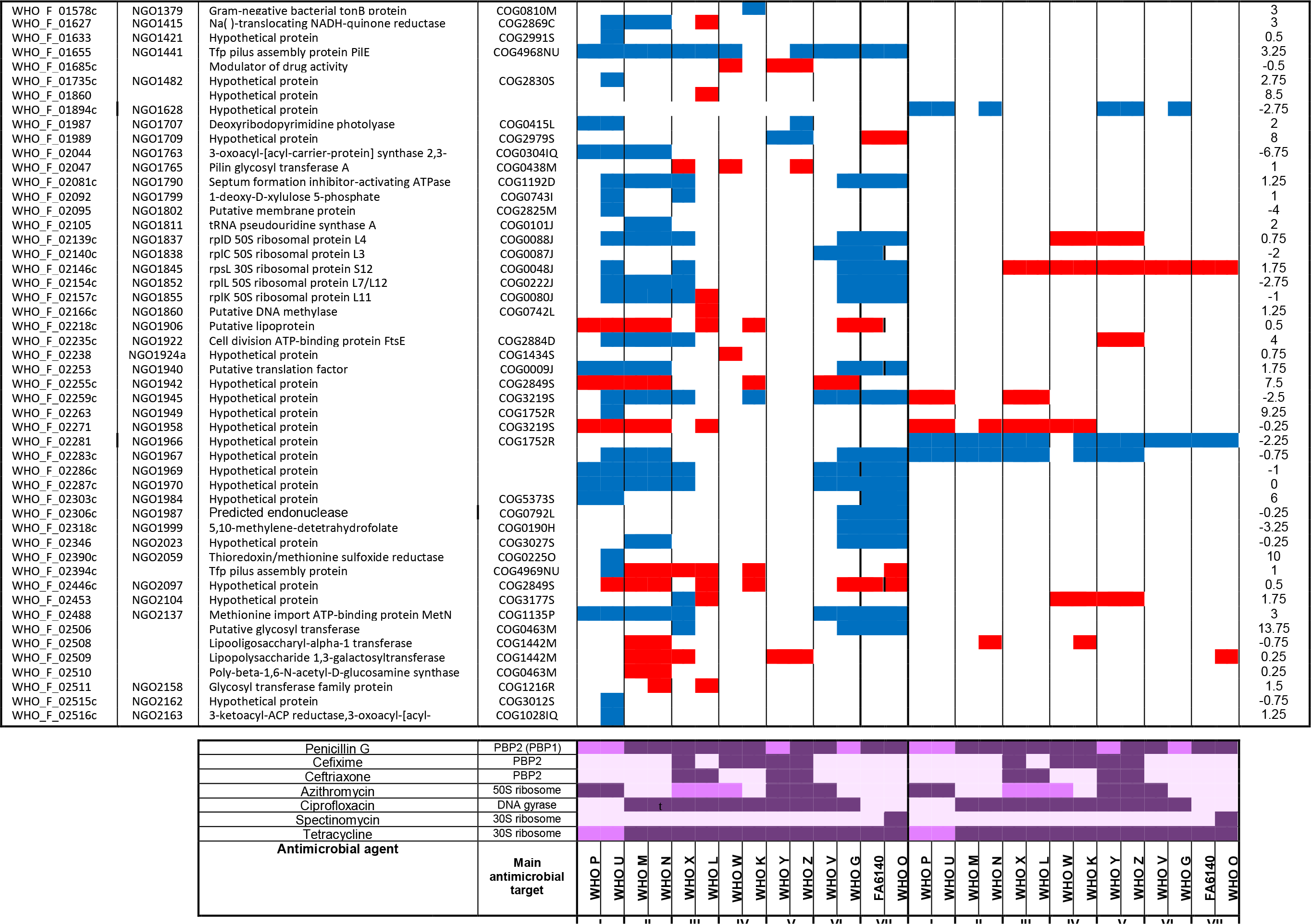

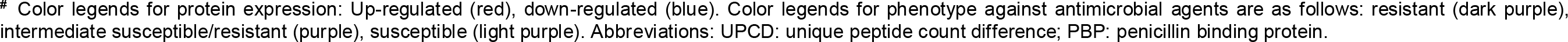

### Proteomic signature of previously verified gonococcal antimicrobial resistance determinants

Our proteomic analysis detected subcomponents of all the five efflux pumps described in *N. gonorrhoeae*, i.e., MtrCDE, MtrF, FarAB, MacAB, and NorM (Table 3). The outer membrane-barrel protein MtrE serves as the channel for the tripartite MtrCDE pump and likely fulfills this same function in the MacAB and FarAB efflux pumps (133, 135). The MtrCDE complex is the most studied efflux pump system in *N. gonorrhoeae*. The multiple transferable resistance (*mtr*) locus contains the *mtrCDE* operon (136) that is negatively regulated by the repressor MtrR (137). Mutations that abrogate *mtrR* activity result in an over-expression of the MtrCDE efflux pump and decreased susceptibility to numerous antimicrobials, e.g. macrolides, penicillins, cephalosporins, and tetracycline (53, 135). AMR mutations in the *mtrR* promoter were previously identified in WHO G, K, M, O, P, V, W, X, Y, and Z and within the *mtrR* gene (G45D or a frame shift mutation resulting in a truncated peptide) in WHO K, L, M, P, and W (50), suggesting an over-expression of the MtrCDE efflux pump. Our proteomic profiling also verified that the levels of MtrE were significantly increased in all isolates with the exception of two strains lacking any type of *mtrR* AMR determinant (WHO U and N; Table 3). Accordingly, MtrE proved to be an effective indicator of expression of the MtrCDE efflux pump. Our findings were further supported by the down-regulation of MtrR in all examined strains except WHO L (Table 3). WHO L is also the only examined strain that contains an *mtr*_120_ mutation, which generates a novel promoter for *mtrCDE* transcription and further enhances the expression of the MtrCDE efflux pump (50, 138). The second efflux system that showed differential expression was the MacAB efflux pump, which can decrease macrolide susceptibility (139). MacA expression varied across the isolates. Expression of the inner membrane component, MacB, was enhanced in the azithromycin resistant strains WHO P and V, but also in the azithromycin susceptible strains WHO N and K, as well as WHO L, which is intermediately susceptible to azithromycin. Interestingly, MacB was the most highly expressed in WHO V, the only strain with high-level azithromycin resistance [MIC>256 µg/mL; due to the 23S rRNA A2059G mutation in all four alleles (50)], which indicates that over-expression of the MacAB efflux pump may contribute to the high MICs of azithromycin and other macrolides in WHO V. The FarA component of the FarAB efflux pump system, which exports long-chain fatty acids and other hydrophobic agents (140), was not over-expressed in any of the examined strains and was instead ubiquitously expressed in seven WHO strains (M, N, K, L, X, Z, and V) and down-regulated in WHO U, O, FA6140. Our proteomic profiling also revealed that the NorM and MtrF efflux pumps, which can decrease the susceptibility to fluoroquinolones and sulfonamides, respectively (141, 142), were not over-expressed in any of the examined strains.

Among other established AMR determinants that were differentially expressed was the major porin of *N. gonorrhoeae*, PorB (143, 144), which was down-regulated in all strains with the exception of WHO G and N (Table 4). This down-regulation suggests reduced import of antimicrobials such as penicillins, cephalosporins and tetracyclines, which can contribute to a decreased antimicrobial susceptibility. Furthermore, the WHO F, G, and N express PorB1a, which is associated with a lack of the AMR determinant *penB* and consequently high-level chromosomally-mediated resistance to penicillins and cephalosporins (127), while all other strains express PorB1b. All WHO strains with PorB1b (*n*=11), except WHO U, contained the AMR determinant *penB*. Consequently, our proteomic data suggest that *penB* may be associated with also a decreased expression of PorB1b in addition to the previously documented decreased penetration through PorB1b, resulting in a decreased susceptibility to several antimicrobials. The expression of penicillin-binding protein 1 (PBP1) was significantly down-regulated in nine out of the twelve WHO strains that possess the *ponA1* resistance determinant, which encodes a L421P amino acid substitution in PBP1 that contributes to high-level chromosomally-mediated penicillin resistance (50). Accordingly, our proteomic data indicate that the PBP1 L421P amino acid alteration, in addition to decreased expression of PBP1, might contribute to high-level chromosomally mediated penicillin resistance. In contrast, PBP2 (the main lethal target for penicillins and cephalosporins) was ubiquitously expressed in 13 of the 14 WHO strains. Similarly, GyrA expression was ubiquitous in 13 of the 14 WHO strains. No association between GyrA expression and the main fluoroquinolone resistance mutations [amino acids S91 and D95 (50)] was identified. Both GyrB and the second fluoroquinolone target, ParC, were over-expressed in the four ciprofloxacin-resistant WHO strains G, N, V and X, which may suggest that these strains upregulate GyrB and ParC to compensate for the mutated main fluoroquinolone target GyrA (Table 3).

### New potential proteomic-derived antimicrobial resistance signatures

In the CE fraction, two hypothetical proteins predicted to localize to the inner membrane, WHO_F_00238c and WHO_F_01226, were down-regulated in all examined strains compared to the antimicrobial-susceptible WHO F strain (Table 4). WHO_F_00238c, which corresponds to NGO0222 in the FA1090 genome, is a small protein with a predicted molecular weight of 8.32 kDa that contains two predicted transmembrane domains but no signal peptide. WHO_F_01226 lacks a homologous protein in FA1090. This is also a small protein (5.39 kDa) with no peptides predicted to be recognized by signal peptidase I or II. In the C fraction, no protein was differentially expressed in all strains, but two cytoplasmic proteins, NGO0597 and NGO0701, were up-regulated in all strains except WHO L. NGO0597 is a nucleoside diphosphate kinase (Ndk; 15.4 KDa) involved in DNA and RNA synthesis (145), regulation of gene transcription (146), and peptide chain elongation during translation (147), all processes that are targets for different antimicrobials. NdK is secreted from *Pseudomonas aeruginosa* (146), *M. tuberculosis* (148), *and Leishmania* (149) to modulate interaction with host cells, block phagosome maturation in macrophages (148, 150), and promote host cell apoptosis and necrosis (151). It remains to be investigated whether the gonococcal Ndk is secreted during infection and whether it may serve as an anti-virulence or antimicrobial target. Finally, we detected two proteins with undefined subcellular localization displaying global differential expression. WHO_F_01139 and WHO_F_01144, which have no homologs in the FA1090 genome, were down-regulated in all strains. Our use of UPCD predicted WHO_F_01139 to localize to the cell envelope. WHO_F_01139 is a putative lipoprotein (16.9 KDa) with a predicted signal peptide II domain. Based on UPCD, in addition to the lack of a predicted signal peptide and the absence of transmembrane domains, we predict the hypothetical protein WHO_F_01144 (7.4 kDa) is cytoplasmic. The impact of these six proteins on AMR is yet to be elucidated; however, our data suggest that they may represent general proteomic markers for gonococcal AMR, a predisposition toward developing or compensating for gonococcal AMR, and/or new antimicrobial targets.

### Phenotypic clustering based on antibiograms and common differentially expressed proteins

To link AMR phenotypes with proteomic signatures, we performed phenotypic clustering of gonococcal strains based on their defined antibiograms (50, 53–55, 152) and common differentially expressed proteins (Tables 4–5). We additionally investigated each protein’s Cluster of Orthologous Genes (COG) annotations and inferred the functional relevance to the observed phenotypes. These analyses generated seven phenotypic clusters that matched between established and proteome-derived AMR signatures (I-VII; Tables 4–6).

Cluster I strains, WHO P and U, exhibit resistance to azithromycin and the majority of up-regulated proteins identified were involved in ribosomal biogenesis: 30S ribosomal proteins S15 and S19; 50S ribosomal proteins L1, L2, and L22; the small GTPase EngA; pseudouridine synthase; RNA helicase; and ribonuclease E. In contrast, proteins involved in cell envelope biogenesis - PilE, LolA, and PglB - were down-regulated in both strains, which may be associated with the strains’ decreased susceptibility to penicillin G.

Cluster II strains, WHO M and N, exhibit resistance to penicillin G, tetracycline, and ciprofloxacin. Up-regulated proteins included DNA repair factors (DnaE, a putative type I-site specific deoxyribonuclease NGO0407, and exonuclease UvrB). Proteins involved in amino acid metabolism (NGO0269, NGO0679); and translation (NGO0803, NGO1870) were also up-regulated, which suggested strains in Cluster II possess compensatory mechanisms for ciprofloxacin and tetracycline resistance, respectively. Hypothetical proteins represented the majority of down-regulated proteins and included WHO_F_00875c, NGO1299, NGO1945, NGO1967, NGO1969, NGO1970, and NGO2089.

Cluster III, IV, and V are comprised of WHO X and L, WHO W and K, and WHO Y and Z, respectively, and exhibit resistance to at least four different antimicrobials, with ciprofloxacin and tetracycline in common (Table 4–5). Three proteins in common between strains in Cluster III and IV were identified. Of these, homoserine dehydrogenase and holo-ACP synthase - involved in amino acid and lipid metabolism, respectively - were down-regulated, while thioredoxin, which is involved in defense against oxidative stress and protein turnover, was up-regulated (153, 154). The NADP quinone reductase (MdaB, modulator of drug activity B) was up-regulated in Cluster IV and V strains. In *E. coli*, MdaB protects against polyketide compound toxicity (155), while overproduction of this protein defends *P. aeruginosa* from oxidative stress (156).

Strains in Cluster VI (WHO V and G) are resistant to penicillin G (WHO G intermediate susceptible), tetracycline and ciprofloxacin. Differentially regulated proteins in this cluster were strikingly similar to Cluster II, with 13 proteins in common (Table 4–6). Seven proteins involved in DNA repair, amino acid metabolism and translation were up-regulated, further strengthening a possible compensatory mechanism for the resistance to ciprofloxacin and tetracycline. Six proteins functioning in coenzyme metabolism (NGO2056) and with unknown functions (NGO1299, NGO1945, NGO1969, NGO1970, NGO2089) were down-regulated (Table 6).

**TABLE 6.**
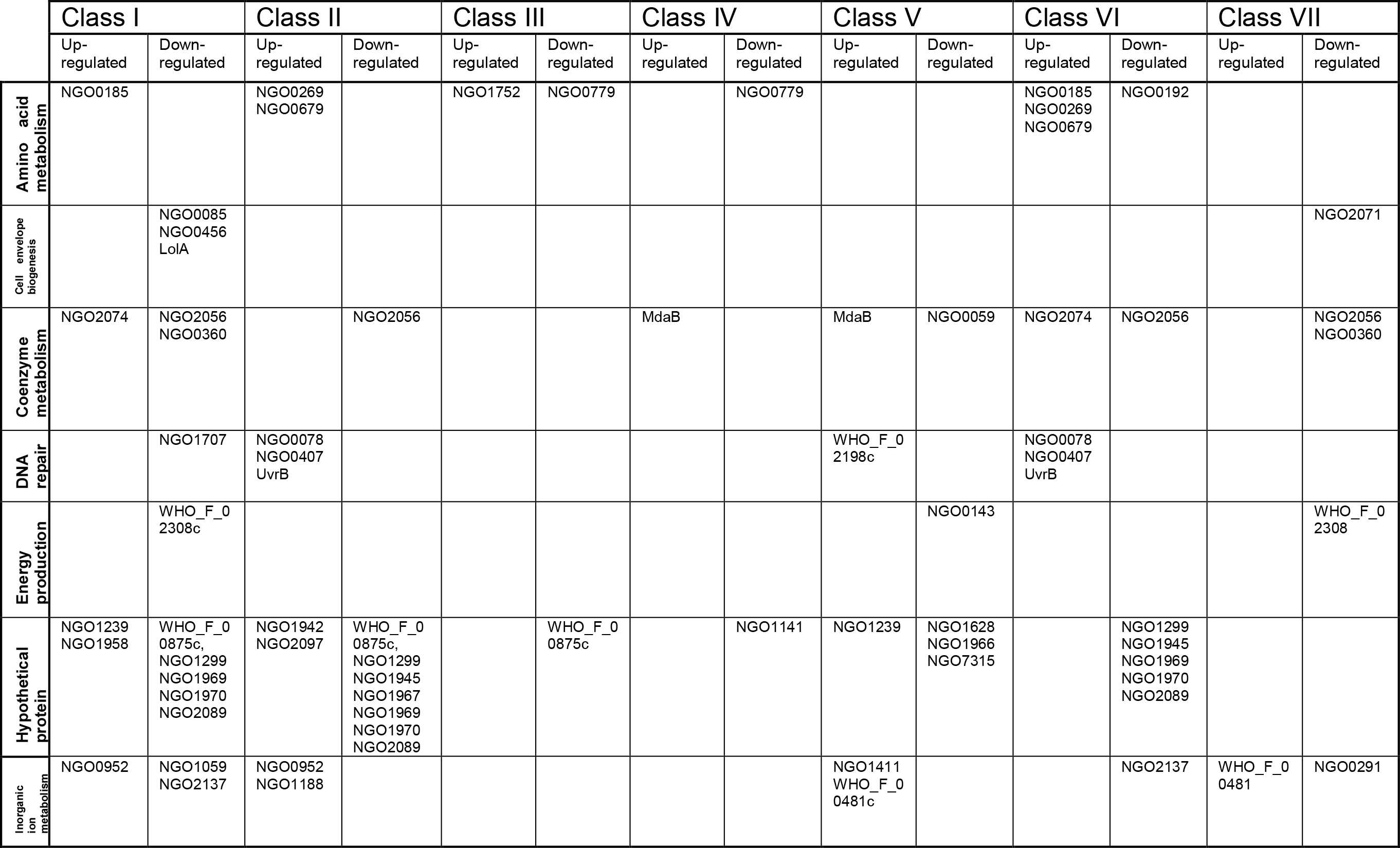
Common differentially expressed proteins in phenotypically clustered *Neisseria gonorrhoeae* strains. Proteomic mining of gonorrhea antigens and AMR.

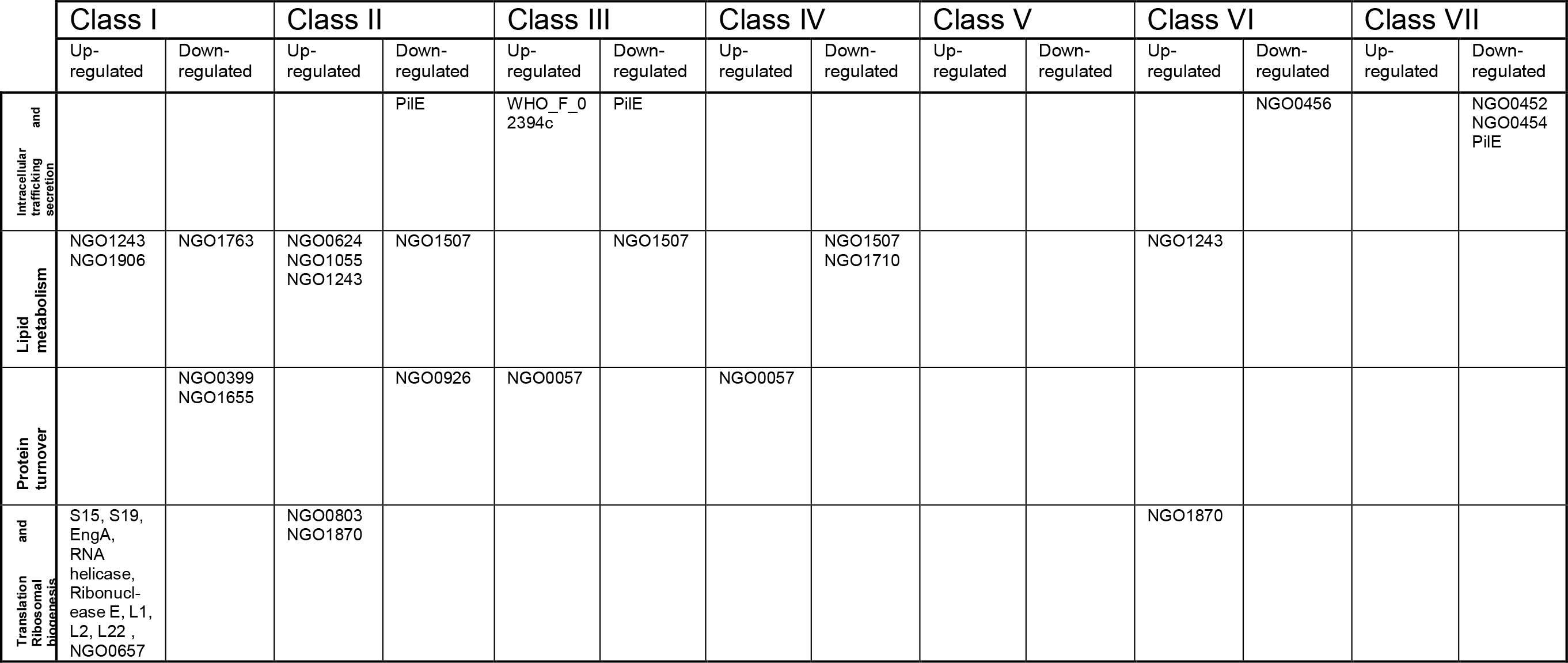

Finally, the cluster VII strains (WHO O and FA6140), displaying resistance to penicillin G and tetracycline, had nine common differentially expressed proteins (Table 6). Among these proteins, two metabolic coenzymes (NGO0360 and NGO2056) and a putative cytochrome *b*561 involved in energy production were down-regulated. This cluster possessed a similar expression profile to strains in Cluster I that are intermediately susceptibility to penicillin G and tetracycline. Finally NGO2017, a putative integral inner membrane protein; NGO0452, a potassium proton/antiporter; PilW; and PilE were also down-regulated in the cluster VII strains (Table 6).

Our proteomic findings elucidate many differentially regulated proteins as potential general proteomic markers for gonococcal AMR, a predisposition toward developing or compensating for gonococcal AMR, and/or new antimicrobial targets, e.g. NGO0222, WHO_F_01226, NGO0597, NGO0701, WHO_F_01139, WHO_F_011144. Deeper analysis of gonococcal proteotypes that relied on AMR-based phenotypic clustering identified additional proteomic markers potentially associated with (or compensating for) AMR in clusters I, II, VI, and VII. Further studies should examine the proteomic profiles of wild type and AMR gonococcal strains during exposure to varying levels of different antimicrobials. In line with this, the expression of eight outer membrane proteins was enhanced in ampicillin resistant *E. coli* strains upon exposure to the minimal inhibitory concentration of ampicillin (33). Additionally, the functional role(s) of the differentially regulated hypothetical proteins potentially involved in gonococcal AMR need to be elucidated, which would help decode the intricate AMR network and promote the design of ways to curb the spread of AMR among *N. gonorrhoeae* strains.

## CONCLUSIONS

The present study provides the first global quantitative proteomic characterization of the 2016 WHO *N. gonorrhoeae* reference strains (50) and FA6140 (51) to identify new vaccine candidates, gain information about expression of previously identified antigens, and enhance our understanding of AMR in *N. gonorrhoeae*. To our knowledge, this is also the largest quantitative proteomics study performed on bacterial sub-proteomes to date. Importantly, nine novel vaccine candidates have been identified, significantly broadening the gonorrhea antigen repertoire. Further, expression of 21 previously verified AMR determinants at the proteome level was investigated and six new proteomic signatures that may be associated with AMR or may indicate a strain’s likelihood of developing or compensating for the physiological consequences of gonococcal AMR. The proteomic signatures we identified may also represent new antimicrobial targets. Expression patterns of antimicrobial targets and AMR determinants provide proteomic signatures that can complement, verify, and enhance our phenotypic-and genetic-derived understanding of gonococcal AMR complexity. Cumulatively, our studies provide a wealth of information regarding gonococcal proteomic profiles and will contribute to ongoing efforts in vaccine/drug development as well as elucidation of AMR mechanisms in *N. gonorrhoeae*.

## ACKNOWLEDGEMENTS

This work was supported by the National Institute of Allergy and Infectious Diseases R01-AI117235 to A.E.S., and the World Health Organization, Geneva, Switzerland (2015) and the Foundation for Medical Research at Örebro University Hospital, Sweden (2016) to M.U. We thank Benjamin I. Baarda for critical reading of this manuscript.

## The abbreviations used are

ACN: Acetonitrile
AGC: Automatic gain control
AMR: Antimicrobial resistance
C: Cytoplasmic
CDC: Centers for Disease Control and Prevention
CE: Cell envelope
COG: Cluster of orthologous genes
cRAP: Common repository of adventitious proteins
FDR: False discovery rate
GCB: Gonococcal base agar
GCBL: Gonococcal base liquid medium
KEGG: Kyoto encyclopedia of genes and genomes
LPS: Lipopolysaccharide
OMV: Outer membrane vesicle
ORF: Open reading frame
WHO: World Health Organization

